# An appraisal of the classic forest succession paradigm with the shade tolerance index

**DOI:** 10.1101/004994

**Authors:** Jean Lienard, Ionut Florescu, Nikolay Strigul

## Abstract

In this paper we revisit the classic theory of forest succession that relates shade tolerance and species replacement and assess its validity to understand patch-mosaic patterns of forested ecosystems of the USA. We introduce a macroscopic parameter called the “shade tolerance index” and compare it to the classic continuum index in southern Wisconsin forests. We exemplify shade tolerance driven succession in White Pine–Eastern Hemlock forests using computer simulations and analyzing approximated chronosequence data from the USDA FIA forest inventory. We describe this parameter across the last 50 years in the ecoregions of mainland USA, and demonstrate that it does not correlate with the usual macroscopic characteristics of stand age, biomass, basal area, and biodiversity measures. We characterize the dynamics of shade tolerance index using transition matrices and delimit geographical areas based on the relevance of shade tolerance to explain forest succession. We conclude that shade tolerance driven succession is linked to climatic variables and can be considered as a primary driving factor of forest dynamics mostly in central-north and northeastern areas in the USA. Overall, the shade tolerance index constitutes a new quantitative approach that can be used to understand and predict succession of forested ecosystems and biogeographic patterns.

## 1 Introduction

Forest succession is a continuous stochastic process that occurs at the level of individual trees and results in the replacement of one tree species by another at the level of forest stands (Horn, 1974, White, 1979, McCarthy, 2001). Understanding the mechanisms underlying forest succession has remained a challenging scientific problem for more than a century (Clements, 1916, Pickett et al., 1987, Huston and Smith, 1987). Existing succession theories are mostly qualitative and represented as conceptual models having substantial limitations in their appraisal and validation (Connell and Slatyer, 1977, Frelich and Reich, 1999, Platt and Connell, 2003). Quantitative forest descriptions are typically based on simple indicators (e.g., time since the last major disturbance) or on macroscopic stand characteristics calculated using forest surveys (e.g. stand age and size structures, tree abundance, biodiversity, and biomass). However, these parameters are only remotely linked with forest succession. The lack of quantitative characteristics capable to quantitatively describe successional patterns in space and time hinders our ability to understand and predict changes of forested ecosystems. Ideally, we would like to have quantitative variables describing forest succession under different disturbance regimes. Such variables can be used to analyze stand dynamics and spatial forest patterns within already existing modeling frameworks, such as the hierarchical patch dynamics concept (Jianguo and Loucks, 1996, Strigul et al., 2012).

The classic succession paradigm has been formulated based on observations of temperate forest patterns in Wisconsin, Michigan and New England (Clements, 1916, Cowles, 1911) and in northern and Central Europe (Sukatschew, 1928). In this type of forest the gap dynamics and shade tolerance driven succession are most noticeable and easy to observe. In a broad range of plant ecology literature, including in major textbooks, shade tolerance is considered as a primary factor underlying forest successional dynamics (Spurr and Barnes, 1980, Walters and Reich, 1996, Smith, 1997). North-American tree species can also independently be classified as early and late successional species based on their life history and physiological traits (Parrish and Bazzaz, 1982, Huston and Smith, 1987). In the classic shade tolerance succession paradigm, species that are shade intolerant and tolerant are analogous to early and late successional species, respectively (Bazzaz, 1979). Replacement of early successional trees by late successional trees is driven by small scale disturbances caused by wind, tree diseases and tree removal (Yamamoto, 2000). Large scale disturbances such as hurricanes, severe forest fire, clearcutting and some rare catastrophic events e.g. volcano eruptions significantly change the successional stage of forest stands by promoting development of early successional species (Bormann and Likens, 1979, Pickett et al., 1989). Particular disturbances often have an intermediate scale and can be placed on a continuous scale of disturbances (Łaska, 2001, Turner, 2005). The interplay of disturbance regimes and successional processes results in complicated and poorly understood multidimensional spatiotemporal patterns (Turner, 2005).

The goal of our research is to develop a quantitative approach that can be used to appraise succession of forested ecosystems. In the present work we introduce a macroscopic stand characteristic called the shade tolerance index defined as a weighted average of shade tolerant trees in a stand. We employ this new variable to describe patch-mosaic properties of the U.S. forests using the Forest Inventory and Analysis (FIA) Program data (http://www.fia.fs.fed.us/). This database has been collected by the U.S. Forest Service since the late 60s by surveying forested ecosystems across all U.S. ecological domains. We compare the shade tolerance index to the continuum index in southern Wisconsin forests using surveys from the FIA database, matching the location of the classic study of Curtis and McIntosh (1951). We also assess the relevance of shade tolerance index to understand forest succession in White Pine – Eastern Hemlock forest of northeastern US. This is done using two approaches: analysis of approximated chronosequence from the FIA database and individual-based computer simulations. We describe the shade tolerance index in mainland USA, and we demonstrate that it does not correlate with other macroscopic characteristics of forest stands. We finally characterize the dynamics of shade tolerance index based on re-sampled plots in the database and we sketch geographical areas based on the importance of shade tolerance as a driving force for forest succession. The appendices include description of the FIA database used in this study, quantitative tables of shade tolerance for North-American trees, and additional statistical results.

## 2 Shade tolerance index and succession dynamics

### 2.1 Shade tolerance and the classic model of succession

Shade tolerance is the ability of a tree to survive and develop under light limited conditions. Overall, tree species can be ranked by their tolerance to different factors affecting tree growth and survival, such as pollutants, heat, drought and nutrient deficiency. Therefore, shade tolerance can be thought of as just one of these tolerance types (Shirley, 1943, Niinemets and Valladares, 2006, Hallik et al., 2009). However, shade tolerance plays a special role in forest development since light is one of the most important factors for tree growth (Zon and Graves, 1911, Shirley, 1943, Baker, 1949). Ecological effects of shade are connected with all aspects of a plant life cycle including growth, reproduction, and mortality. This was noticed in numerous publications by both professional foresters and amateur naturalists. In particular, Pliny the Elder described some ecological effects of shade more than 2000 years ago (Pliny the Elder, 1855, Chapter 18). Henry David Thoreau wrote in his famous 1860 essay on forest tree succession: “The shade of a dense pine wood, is more unfavorable to the springing up of pines of the same species than of oaks within it, though the former may come up abundantly when the pines are cut, if there chance to be sound seed in the ground” (Thoreau, 1992). The shade tolerance concept based on empirical data was developed in the XIX^th^ century and substantially influenced further development of plant succession theory in XX^th^ century (Cowles, 1911, Clements, 1916, Connell and Slatyer, 1977). In particular, as early as 1852, German forester Heyer (1852) has published this conceptual framework and a shade tolerance ranking table of European trees. Extended studies of the late 1800s to early 1900s resulted in hundreds of research papers and several books providing synthesis and shade tolerance tables for tree species in the northern Hemisphere (Morozov, 1908, 1912, Zon and Graves, 1911, Sukatschew, 1928).

The shade tolerance tables classify tree species by their shade tolerance patterns along a linear scale (Zon and Graves, 1911, Baker, 1949, Valladares and Niinemets, 2008). A typical shade tolerance classification consists of 5 uniformly distributed values: very intolerant, intolerant, intermediate, tolerant and very tolerant trees. The earlier classifications have been based on empirical observations of tree species growth under light limiting conditions, including leaf density, rate of death of the bottom tree branches shadowed by the upper tree branches (self-pruning), self-thinning of forest stands, density of trees in the understory and tree mortality (Morozov, 1908, 1912, Zon and Graves, 1911). Early experimental studies examined physiological and morphological mechanisms underlying shade tolerance, including differences in leaf morphological structure and photosynthetic activity to discover mechanisms of shade tolerance (Morozov, 1908, 1912, Zon and Graves, 1911). The modern species-specific shade tolerance tables have the same empirical character, but have been developed using the survey method. This is where teams of professional experts independently evaluate different plant species with respect to their shade tolerance and the final ranking is represented as a statistics of these independent evaluations (Baker, 1949, Humbert et al., 2007). In particular, the shade tolerance tables for the US forests proposed by Zon and Graves (1911) have been based on empirical observations and designed to be analogous with European trees, for which several different classifications had been previously established. These tables were revised for the first time 30 years later by Baker (1949), who employed summarized opinions of 55 foresters as well as published studies. The next substantial revision came after another 50 years, when Humbert et al. (2007) provided the shade tolerance values not only for tree species but also for shrub, herbaceous, bryophyte and lichen species. These consequent revisions obtained with the survey methods expanded the range of species covered and have overall confirmed the earlier ranking of the major US tree species proposed by Zon and Graves (1911). These existing shade tolerance rankings are species-level expectations of individual-level random variables, and it is broadly recognized that there exists substantial interspecific variability of shade tolerance depending on the environmental conditions and tree age (Zon and Graves, 1911, Baker, 1949, Humbert et al., 2007, Kunstler et al., 2009, Valladares and Niinemets, 2008). At the same time numerous quantitative studies based on the species-level shade tolerance values demonstrate that shade tolerance is one of the major physiological traits correlated with other traits such as understory mortality and growth (Kobe et al., 1995, Walters and Reich, 1996, Lin et al., 2001, Valladares and Niinemets, 2008), photosynthetic and allocation patterns (Kitajima, 1994), tree morphology (Messier et al., 1999), crown ratio (Lorimer, 1983), root distribution (Gale and Grigal, 1987) and seed size (Hewitt, 1998).

The concept of shade tolerance is naturally linked with the concepts of forest succession and disturbance. The classic successional model employs shade tolerance to explain the following transitions in canopy tree composition after a major disturbance (Canham, 1988, Smith, 1997, McCarthy, 2001):

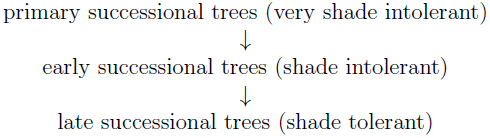

This conceptual model historically states that the initial canopy is formed mostly by primary and early successional species that are shade intolerant but tolerant to numerous adverse factors affecting tree growth in the absence of the microclimate created by forest canopy, for example water limitation, extreme temperatures, ice damage (Morozov, 1908, Sukatschew, 1928, Shugart, 1984, Oliver and Larson, 1996). After the canopy is closed, forest succession is driven by individual tree mortality and small disturbances that develop relatively small gaps approximately equal to one average crown area which facilitate recruitment of shade tolerant understory trees (Runkle, 1982, Pickett et al., 1987, Frelich and Martin, 1988, Canham, 1988, Smith, 1997). The conceptual model of the forest gap dynamics (White, 1979, McCarthy, 2001) includes two processes: 1) the development of forest gaps in the canopy after the death of large trees, and 2) recruitment of understory trees into the canopy. The gap facilitates recruitment of understory trees which are mostly shade tolerant (Yamamoto, 2000). Intermediate and large scale disturbances can create large canopy openings and promote development and recruitment of faster growing early successional tree species, which are mostly shade intolerant (Łaska, 2001, Platt and Connell, 2003).

### 2.2 The shade tolerance index

According to the classic paradigm described in the previous section, the proportion of shade tolerant versus intolerant trees is linked to the forest succession stage. We propose a quantitative parameter, the *shade tolerance index*, *δ*, to characterize *stand successional stages* as follows:

1. Shade tolerance of every *tree species* is quantified by a number from an interval *ρ* = [0, 1] where the range spans very intolerant to tolerant species. We will call the number *ρ* the *shade tolerance rank of a tree species*.
2. *δ*, the shade tolerance index of the stand is defined as a weighted sum of the species abundance on their shade tolerance ranks defined as:

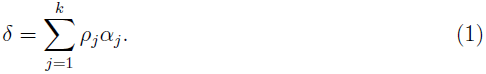

In this formula *α_j_* is a measure of relative abundance of a species *j* in the stand, *ρ_j_* is a shade tolerance rank of the species *j*, and the index *j* runs through all *k* tree species present in the stand.

The relative abundance parameter *α_j_* is estimated using the formula:

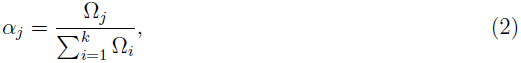

where Ω*_j_* is a measure of abundance of the tree species in the stand. This measure Ω*_j_* may be calculated using all the trees in the stand or only for some particular group of trees, for example the canopy or the understory trees.

The relative abundance measure *α_j_*, is a number in the interval [0, 1] and obviously 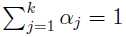 regardless of the choice for the measure Ω*_j_*. Since all *ρ_i_* are in [0, 1], the shade tolerance index *δ*, is also a number from [0, 1]. Specifically, *δ* is equal to 0 if all the trees in the stand are very shade intolerant and equal to 1 if all trees are shade tolerant. In Appendix 1, we address the technical points of the practical calculation of Ω*_j_* as well as *ρ_j_*. We also investigate the robustness of the *shade tolerance index of the stand δ* with respect to the way in which Ω*_j_* and *ρ_j_* are defined.

The shade tolerance index can be calculated at the level of individual trees with *ρ* as a function of a tree life history, size, age and environmental factors, however, given the lack of such information, we employ the average tree species values according to the shade tolerance ranking tables. Specifically, we quantify the species as following: very intolerant = 0, intolerant = 0.25, intermediate = 0.5, tolerant = 0.75, and very tolerant = 1 (Appendix 1).

### 2.3 Comparison with the forest continuum index

The classic forest succession paradigm was historically developed under a strong influence of the studies conducted in the Lake States (MI, WI and MN). In particular, both Clements (Clements, 1916) and Cowles (Cowles, 1911), the founders of this paradigm, have done their field research in this area. Later, Curtis and McIntosh (1951) have developed the classic continuum index to describe successional patterns in southern Wisconsin. This landmark paper was an important step in the development of forest succession theory that influenced numerous studies and theoretical developments, in particular the continuum hypothesis of vegetation (McIntosh, 1967, Nakamura, 1985), gradient analysis of vegetation (Whittaker, 1967) and ordination of plant communities (Bray and Curtis, 1957, Austin, 1977, 1985). We anticipate that the shade tolerance index is capable of capturing successional patterns, and we compare these indexes in southern Wisconsin.

Curtis and McIntosh (1951) studied successional patterns in a collection of 95 selected forest stands. The authors selected this snapshot of the forest stands at different successional stages to characterize the continuous transition from early successional stages to the climax stage. The importance values corresponding to tree species have been computed based on respective species dominance within the stands. Using tree dominance data, these stands were positioned along the continuum line representing a forest succession sequence. According to this continuum index axis, Curtis and McIntosh (1951) assigned numerical values to the tree species within the stands called the climax adaptation numbers. The authors have also demonstrated that the constructed continuum index is related to soil factors such as exchangeable calcium concentration and soil acidity.

Curtis and McIntosh (1951) did not employ the concepts of shade tolerance and gap dynamics to derive the climax adaptation numbers. However, they indicated that the calculated climax adaptation numbers are parallel to the shade tolerance ranking used in forestry (Curtis and McIntosh, 1951, pp. 489-490). Our mechanistically-based approach of obtaining the successional axis is substantially different from the empirical-based approach employed by Curtis and McIntosh (1951). In particular, the shade tolerance index is based on independently established shade tolerance rankings, while the continuum index relies solely on the composition of sampled plots.

The original data used by Curtis and McIntosh (1951) is not available, and we reproduced a similar analysis using the FIA data. We have calculated the same statistical characteristics using a sample of about 7000 FIA plots on mesic soils corresponding to the same geographic area as the original article did by using Figure 1 in Curtis and McIntosh (1951) as a reference (see Figure 1 in the Appendix 2). The original study took into account only forest stands that were: 1) “natural forests (i.e. not artificially planted)” of a minimum size of 40 acres, 2) “free from disturbances in the form of fire, grazing or excessive cutting”, and 3) “upland land forms on which run-off water never accumulate” (Curtis and McIntosh, 1951, p. 480). Similarly to the original study, we restricted the analysis to mesic soils (criterion 3). However, in contrast with Curtis and McIntosh (1951) we analyzed all forested plots, including planted forests (criterion 1), and we did not filter out plots based on environmental disturbances (criterion 2).

**Figure 1:**
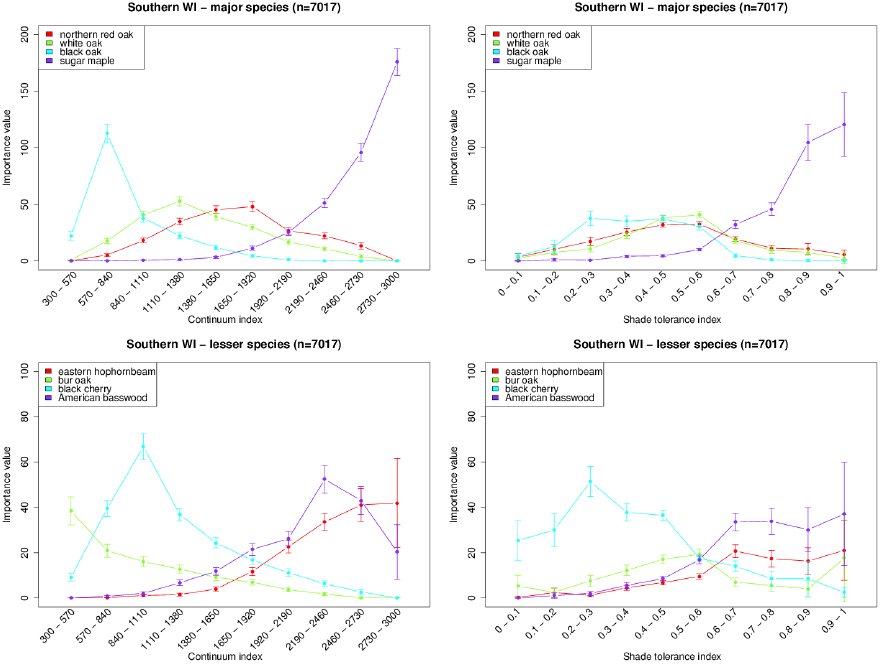
Comparison of the continuum index (left) and shade tolerance index (right) in southern Wisconsin. Similarly to the original study (Curtis and McIntosh, 1951), the species were split in two groups: major (top) and lesser (bottom) species. Bars indicate the standard error of the mean.

Our extension of Curtis and McIntosh (1951) analysis to a wider range of plots resulted in a more diverse species composition. Indeed, only 21 species were considered in the original study, while the actual database plots that we used contain 84 species. To be able to compute the original importance values (Curtis and McIntosh, 1951), we only consider the trees belonging to the 21 species in the original work when computing both the continuum and well as the shade tolerance index. This modification led to a smaller magnitude of the importance value, but overall our results are in good agreement with the original results (Figure 1 vs Figures 5,6,7 in Curtis and McIntosh, 1951). Our results are also in agreement with the work of Rogers et al. (2008) who re-sampled the same sites as Curtis and McIntosh (1951) some 50 years later. In particular, we notice that the relative importance value of red oak (*Quercus rubra*), black oak (*Quercus velutina*) and secondarily white oak (*Quercus alba*) have decreased compared to sugar maple (*Acer saccharum*).

Overall, the successional pattern observed by Curtis and McIntosh (1951) did not change substantially over the several decades, and the original plot sampling restrictions do not substantially affect the observed patterns. At the same time, we have been able to observe successional patterns on a broader range of the continuum index than Curtis and McIntosh (1951) (continuum index changes from 300 to 3000 in Figure 1 and only from 632 to 2650 in the original paper). There are some remarkable successional patterns observed on these additional intervals of the continuum index scale ([300 — 700] and [2600 — 3000]). In particular, black oak has an obvious peak of its importance value at about 700 (Figure 1), that was not observed in the original study and the sugar maple’s importance value monotonically increases along the continuum index. There is a good qualitative correspondence of the species ordination between the continuum index axis and the shade tolerance axis (Figure 1), demonstrating the ability of the shade tolerance index to describe forest succession. Other traditional indicators such as biomass, basal area, biodiversity or stand age do not allow to discriminate between species (Figures 2 and 3 in Appendix 2).

### 2.4 Successional dynamics

The forest stand dynamics theory states that after a major disturbance stand development follows four consecutive stages: initiation, stem exclusion, understory reinitiation, and old-growth (Oliver and Larson, 1996, Smith, 1997). The *stand initiation stage* marks the onset of succession by regeneration of open space from seed, sprouts and advance regeneration, and lasts until the canopy closes. Different disturbances leave various types of biological legacies providing highly variable initial species composition (Franklin et al., 2007, Swanson et al., 2010). In the second stage of *stem exclusion*, the light-driven competition becomes the major determinant of survival, resulting in a domination of fast-growing early successional species. The third stage, *understory reinitiation*, is characterized by the selective recruitment of understory trees in the canopy through gap-dynamics. The final stage, the *old-growth* corresponds to the climax state of the forest, where the species composition is stable.

The shade tolerance index will be used to interpret and quantitatively measure these dynamics. Figure 2 summarizes the conceptual model of shade tolerance driven successional dynamics after a major disturbance. From the definition of the shade tolerance index, we expect an initial decrease of the index value during the reinitiation and stem exclusion stages. This is due to the higher mortality of shade tolerant species that are less resistant to the large number of unfavorable environmental factors, as well as to their exclusion by faster growing shade intolerant species. During the understory reinitiation stage, the shade tolerance index will monotonically increase due to recruitment of shade tolerant species in the canopy. In the old community stage we expect the shade tolerance index to remain stable in the absence of intermediate and large scale disturbances. This conceptual model is in very good agrement with our results, including analysis of the White Pine – Eastern Hemlock succession presented below and data mining of FIA dataset (Section 3.2 and Appendix 3).

**Figure 2:**
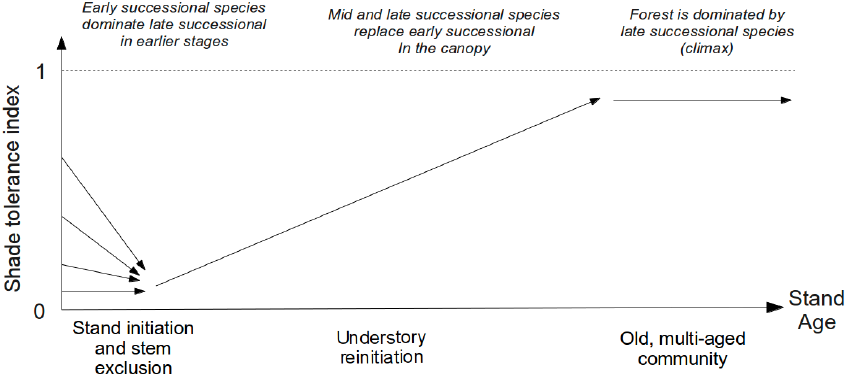
Conceptual model of shade tolerance index dynamics after a major disturbance. The initial proportion of seedling of shade tolerant and intolerant species can be very different due to the nature of the disturbance and biological legacies. Shade intolerant species outgrown shade tolerant species during the stem exclusion stage of the forest stand development, leading to the index decrease. Shade tolerant species gain an edge over shade intolerant species via canopy gap dynamics, resulting in the index increase during the understory reinitiation stage, and its further stabilization as the forest approaches the old growth stage.

We assess the conceptual model of Figure 2 by considering a two-species system consisting of shade intolerant White Pine and shade tolerant Eastern Hemlock. This system embeds the classic shade tolerance paradigm as it demonstrates the shade tolerance trade-off, and at the same time, it plays an important role in the successional dynamics of the temperate forest in the Eastern part of the United States (Bromley, 1935, Nichols, 1935, Burns and Honkala, 1990). We analyze the predictions of a forest gap model (Strigul et al., 2008), and compare them to the approximated chronosequence extracted from the FIA database, in order to examine the shade tolerance dynamics on this bicultural system.

Traditionally, individual-based gap dynamics models are employed to quantitatively predict stand development after major disturbances (Shugart, 1984, Pacala et al., 1993, Bugmann, 2001). Gap models are parameterized by individual-tree data and can be used as a tool to simulate shade tolerance driven succession and statistical characteristics of canopy recruitment (Acevedo et al., 1995, Dubé et al., 2001). The White Pine – Eastern Hemlock system was previously simulated using the crown plastic version of the SORTIE model (Strigul et al., 2008, Figure 12, p. 535), and this model is employed now to assess the temporal dynamics of shade tolerance index. In this model, the shade tolerance trade-off is simplified and limited only to differences in tree growth and mortality, and does not involve interspecific differences in the seed dispersion and reproduction strategy. In particular, both species have the same initial number of seedlings and new seedlings emerge every year in the same quantity. This simulated scenario of the forest succession has been started after a major disturbance, and Figure 3 shows changes of shade tolerance index. The successional dynamics are essentially driven by the shade tolerance trade-off: White Pine is the large tree that grows faster in the full light, but has a higher mortality in the understory, while Eastern Hemlock grows slower but survives in the light limited conditions. The canopy gaps development is randomly driven by individual tree mortality that provides lottery recruitment of understory trees, and this mechanism is sufficient to obtain the early decrease followed by a progressive increase of shade tolerance postulated by the conceptual model of Figure 2.

**Figure 3:**
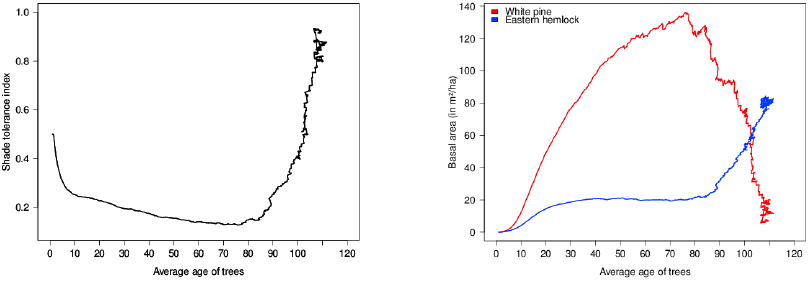
Computer simulation of White Pine - Eastern Hemlock forest stand. Shade tolerance index as a function of the implied stand age (left) and cumulative basal area of White Pine and Eastern Hemlock (right) as a function of the implied stand age. The simulation is conducted using the crown plastic SORTIE model following Strigul et al. (2008) for one hectare and 1000 years, the parameter values are similar to the original paper except the mortality of both species was reduced to 5%.

We compare the results of computer simulations with the statistical analysis of White Pine – Eastern Hemlock forest stands from the FIA database (Figure 4). We consider here approximate chronosequences, as the plots are ordinated relatively to the average age of trees (see Appendix 1), the time since last disturbance being not available. The approximated chronosequence based on the FIA data can be used for model validation in case forest succession data obtained at one particular location are not available (Purves et al., 2008). The stands are observed throughout northeastern parts of the US. We isolated all plots in the database with more than 75% of cumulative basal area composed by these two species, resulting in a pool of 1375 plots (see Figure 1 in Appendix 3 for their locations). The comparison of Figures 4 and 3 demonstrates striking qualitative similarities between computer simulations and the chronosequence. Specifically, the initial distribution of seedling shade tolerance index decreases in the early years as the faster growing pioneering species start to dominate the canopy. These then reach high values as shade tolerant species eventually dominate the early successional species. Similarly, the cumulative basal areas show the same pattern of early dominance of White Pine before Eastern Hemlock take over in old stands. These results are in agreement with the conceptual model (Figure 2).

**Figure 4:**
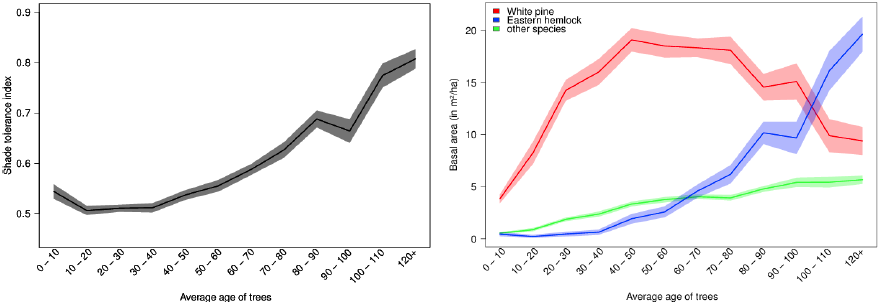
Approximated chronosequence of White Pine – Eastern Hemlock forest stands in the northeastern US. Analyzed data consist of 1375 USDA FIA plots, where these species account for more than 75% of the total basal area (Appendix 3). Shade tolerance index (left) and cumulative basal area of White Pine and Eastern Hemlock (right) as a function of the implied stand age (Appendix 1). The shaded areas represent the standard error of the mean. The last bin “120+” includes 3% of the plots.

## 3 Spatiotemporal patterns of US forests

In this section we analyze forest stand mosaic in mainland US using the FIA dataset. The goal is to understand statistical relationships between forest characteristics and patch mosaic patterns related to shade tolerance. We first conduct a descriptive statistical analysis of shade tolerance patterns, and compare them to indicators of implied stand age, diversity, biomass and basal area. We then study shade tolerance dynamics by grouping all permanent plots according to the Bailey’s ecoregions classification, which relates climatic factors and species compositions (c.f. Bailey, 1995, see Figure 1 in Appendix 1 for a map of the ecoregions). Our analysis allows us to delimit areas and conditions in which shade tolerance is a major driver for succession.

### 3.1 Descriptive statistics and correlations

The stand-level characteristics display different spatial patterns in their distribution across regions (Figure 5). To further study the spatial distribution of the characteristics, we classified plots according to geographically defined areas sharing common ecological properties. We employed Bailey’s province subdivisions because they are fine enough to discriminate between major plan formations, yet large enough to allow for the statistical description of forest patch mosaic (see Figure 1 in Appendix 1). Descriptive statistics further confirm important disparities in the various indicator’s distributions across provinces (Appendix 1).

**Figure 5:**
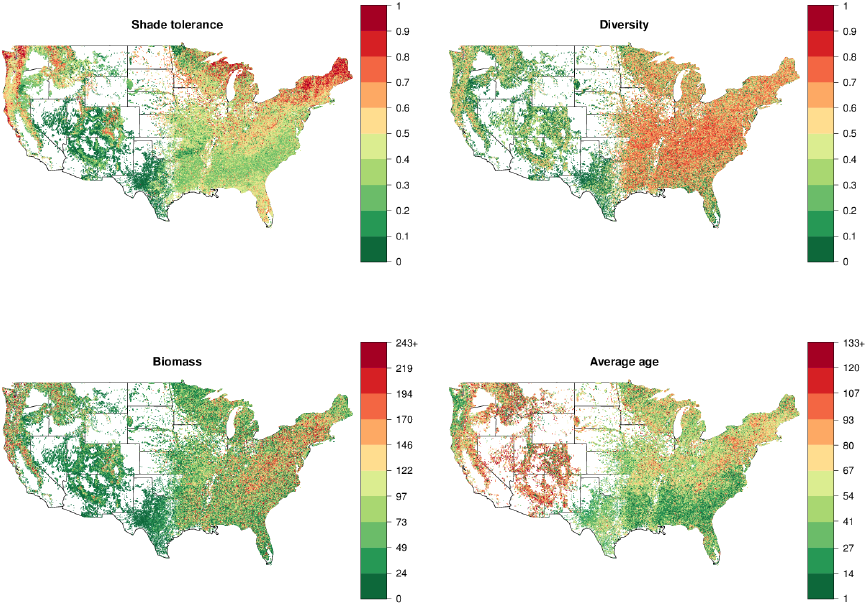
The stand-level characteristics of plots for all years demonstrate very heterogeneous forest types in the US, with no obvious common distribution pattern between indicators. Maps of subsets of plots from different decades are similar and can be found in Appendix 1.

The overall correlation pattern (Figure 6) indicates that the shade tolerance index does not significantly correlate with any other macroscopic characteristics employed in this paper. However, some of the other measures are correlated: biomass with basal area (confirming the previous study of Strigul et al., 2012), and Gini-Simpson diversity with species richness. Correlation matrices have been further calculated separately for all provinces in mainland USA (Figure 5 in Appendix 4) and all inventory years with more than 500 plots recorded from 1968 to 2012 (Figure 4 in Appendix 4). Correlations between different variables were virtually identical for all the inventory years and different ecoregions. This result is similar to what we obtained by analyzing another dataset for Eastern Canada forests (Lienard et al., 2014). The fact that different stand characteristics share a common structure is surprising given the very high variability across North-American forests in terms of species composition, disturbance regimes and land-use history. This stable correlation structure allows to apply multi-variate statistics to define uncorrelated axes describing variability of forest stands through a principal component analysis (Lienard et al., 2014), where the shade tolerance index matches one of the principal component axes. The fact that shade tolerance index has been repeatedly shown to be uncorrelated with other macroscopic characteristics demonstrates its usefulness in the statistical description of the mosaic of forest patches.

**Figure 6:**
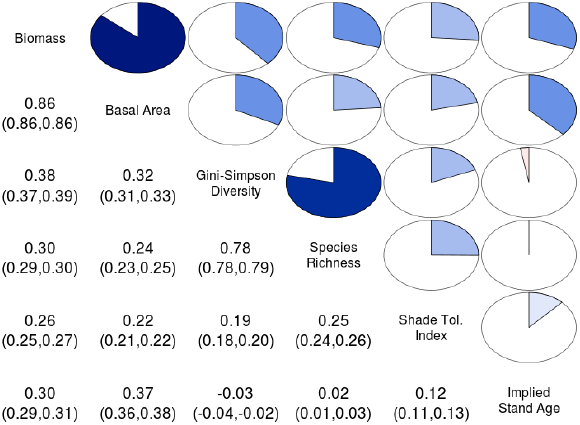
Stand characteristics correlation matrix based on the whole database; the numbers below the diagonal are the Pearsons coefficients (with 95% confidence intervals) and the circles above the diagonal provide visual indications of the correlations. Breakdowns for different ecoregions and different years are presented in Appendix 4 (Figures 5 and 4).

### 3.2 Ecoregion classification

To understand the shade tolerance dynamics, we computed and analyzed the transition matrices for shade tolerance index, by employing an original Bayesian approach (Lienard et al., 2014, see also Appendix 5). We discretized the continuous [0, 1] range of shade tolerance index into 10 even states, the first state being [0, 0.1], and the tenth being [0.9, 1] and estimated a 3-year transition probability matrix in each province. Due to the stochastic nature of Gibbs sampling (Appendix 5), transition matrices computed from a very small number of measurements are meaningless. In this study we consider the transition matrices obtained in provinces with more than 150 remeasured plots (Figure 7).

**Figure 7:**
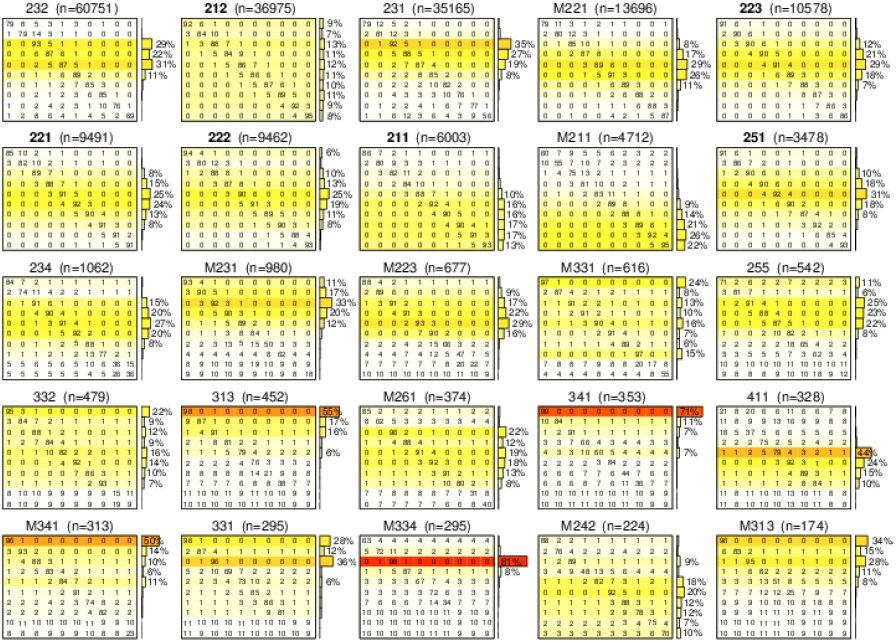
3-year transition matrices of the shade tolerance index for all provinces with n > 150 re-sampled plots. M_i,j_, the value located in row i and column j of a given matrix M, is the probability in percent of transition from state i to state j after 3 years. At the right of the matrices, bar graphs show the overall distribution of plots along the states in the provinces. For easier reading, the color coding for these distributions are also reported in the transition matrices.

The complexity of forest disturbance regimes leads to the non monotonic successional dynamics within stands, which is reflected in the non-zero coefficients of transition matrices (Figure 7). Intermediate and large scale disturbances can promote development of shade tolerant or intolerant trees depending on stand conditions. These disturbances thus cause nonmonotonic behavior (such as sudden jumps and drops) of the shade tolerance index. In particular, in the case of a substantial part of the shade intolerant canopy trees being destroyed and the well-developed shade tolerant understory trees recruited into the canopy, we will observe the jump of the index. In another scenario, an intermediate or large disturbance of shade tolerant tree canopy with no developed understory will facilitate development of shade intolerant species and lead to a drop in the index. These transitions are captured in the matrices of Figure 7.

Ecoregions with succession driven by shade tolerance should have transition probabilities in accordance with the conceptual model of Figure 2, which predicts three different phases of shade tolerance dynamics: (a) a decrease in the early stages, followed by (b) an increase and eventually (c) a stabilization. The matrices of Figure 7 capture all forest transitions disregarding the implied stand age. It is possible to study specifically the initial dynamics of shade tolerance index by computing transition matrices on the subset of plots with a stand age less than 20 years. These matrices (Figure 2 in Appendix 5) demonstrate that a decreasing pattern in the early stages is quite common in the different ecoregions and that the shade tolerance index varies substantially within young forest stands in agreement with the conceptual model. The observed compositions of earlier successional forests are also in agreement with the stand dynamics theory stating that both earlier and late successional species occur during the stand initiation (Swanson et al., 2010, Smith, 1997, Chapter 7, Figure 7.1). However, the limited number of observed transitions of very young stands prevents this analysis to be applied on most US ecoregions. To study systematically the applicability of the shade tolerance succession paradigm to different ecoregions, we further derived two quantitative criteria based on (a) distributions of the shade tolerance index in all plots, and (b) transition matrices in all re-sampled plots (Figure 7).

First, the shade tolerance index should take substantially different values depending on stand age according to the conceptual model of Section 2.4. Indeed, projecting this model on forest stand mosaic within an ecoregion, we anticipate that stands with all possible shade tolerance index will occur, from low values (for plots in the early stages) to high values (for old-growth plots). Quantitatively, this disparity can be assessed by characterizing the shape of the shade tolerance distribution in the forest inventory dataset. In particular, the limited width of the distribution indicates that the shade tolerance is not the primary factor in succession within a given ecoregion. We use the width encompassing 95% of the shade tolerance index as the first criterion for ecoregion classification (Figure 8). The restriction of the distribution width to 95% removes outliers existing in the database that can be seen in distributions of all stand characteristics presented in Appendix 1.

**Figure 8:**
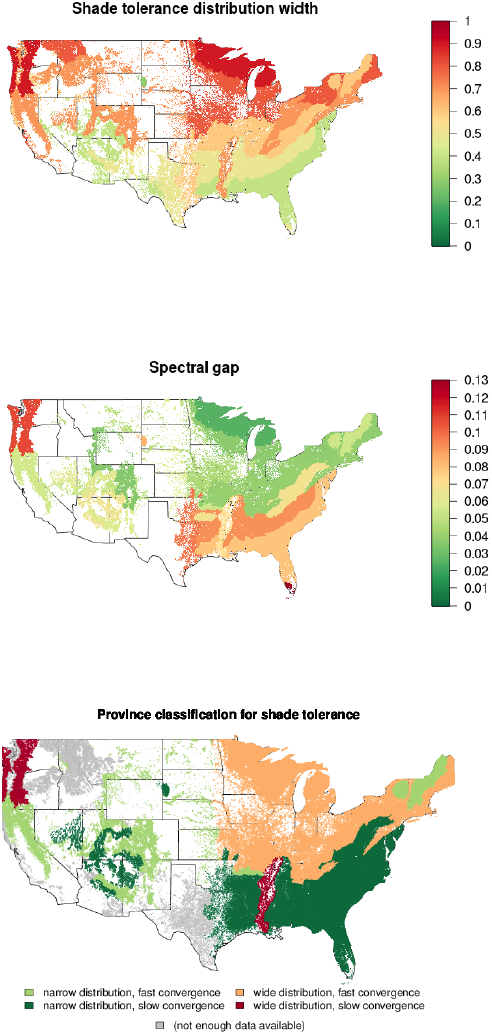
Classification of ecoregions based on shade tolerance distribution width and spectral gap. Top: the disparity of shade tolerance is measured as the interval width containing 95% of the shade tolerance index. Middle: the spectral gap of the transition matrices quantifies the convergence rate after a disturbance, where low values mean that one plot is quick to reach equilibrium after the occurrence of a random disturbance. Bottom: classification of ecoregions. Provinces where forest succession is primarily driven by shade tolerance exhibit a wide distribution of shade tolerance index as well as a fast convergence toward shade tolerance equilibrium.

Second, we expect that shade tolerance should be quicker to reach its equilibrium after a perturbation in the case when shade tolerance is the major driver for succession than in the case when shade tolerance is a secondary factor. To assess this, we compute the spectral gap as a global measure characterizing the average time needed by a plot to reach the shade tolerance equilibrium after a random perturbation. Formally, this metric is defined as 1 *− λ*_2_, where *λ*_2_ is the second highest eigenvalue of the transition matrix (Levin et al., 2009). For a province, high values of this metric mean that the time needed to reach shade tolerance equilibrium is long, which is interpreted as a weak shade tolerance driver for succession. We computed this metric only for provinces for which we can have a reliable estimate of the transition matrices (Figure 8 in main text and Table 1 in Appendix 5).

In the ecoregion classification, we characterize the shade tolerance distributions as either “wide” or “narrow” according to their value compared to the median distribution width (Table 1 in Appendix 5). Similarly, we distinguish between “fast” and “slow” convergence toward equilibrium based on the relative spectral gap value compared to the median spectral gap. We were thus able to delimit different kinds of ecoregions based on their shade tolerance dynamics (Figure 8).

We can distinguish three main types of ecoregion. First, some ecoregions are clearly driven primarily by environmental factors not related to light, and thus do not exhibit different shade tolerance across plots. These regions correspond to the “narrow” shade tolerance distributions in Figure 8 (e.g. deserts in 321, 315 and similar). Second, some ecoregions have succession only driven in a minor way by shade tolerance, with other factors being more important. These regions are characterized by a “slow” convergence toward shade tolerance equilibrium after a disturbance in Figure 8 (e.g. more fire resilient species in fire-driven environments such as 231, 232 and 411). Third, ecoregions that associate both (a) a spatial mosaic of patches with different shade tolerance index and (b) a fast convergence back to shade tolerance equilibrium after a random perturbation, constitute the most suitable provinces to embed the shade tolerance based theory of succession (mostly in north-central and northeastern parts of the USA). Overall, in Figure 8, the succession is orange plots are driven mainly by shade tolerance; red and dark green plots are driven secondarily by shade tolerance; light green plots are driven mostly by other tolerances.

We have applied our shade tolerance classification at the level of the provinces (Figure 8), however the classification provide clear distinctions at higher levels of the biogeographic hierarchy, at the division and domain levels (Figure 2 in Appendix 4; Bailey, 1995). This is an impressive evidence that shade tolerance driven succession is strictly linked with large scale climatic factors and biogeography. Indeed, the biogeographic division of the US forests into domains, divisions and provinces is based on different principles than the shade tolerance based classification (Bailey, 1995, 2004). The shade tolerance classification classification also explains some relationships between spatial distributions of different stand-level characteristics presented in Figure 5. One immediate observation is the clear transitional patterns within the Humid Temperate Domain (200) of Eastern US. Indeed, forests in the Warm and Hot Continental Divisions (210, 220) are driven by shade tolerance succession (Figure 8) and consist of highly diverse forest stand mosaics with respect to biomass, biodiversity and stand age distributions (Figure 5, Appendix 1). On the other hand, forests in the Subtropical Division (230) are not driven by shade tolerance succession and consist mostly of shade intolerant young stands differing in biomass (Figure 5). Lower Mississippi Riverine Forest Province (234) is an obvious exception in this division, as this truly unique province is located in Mississippi River floodplain and consists of low forested terraces and swamps (Bailey, 1995).

Mountain provinces are typically distinct in their classification from surrounding provinces. In particular, M211 (Adirondack - New England Mixed Forest - Coniferous Forest - Alpine Meadow Province) and M221 (Central Appalachian Broadleaf Forest - Coniferous Forest - Meadow Province) are classified as not being driven by shade tolerance whereas other north-eastern ecoregions are classified as being driven by shade tolerance. The vegetation of these two mountain areas is specifically determined by vertical zonation Bailey (1995), where very close patches may be affected by very different climatic conditions, indicating the prevalence of other tolerances over shade tolerance. In addition, mountain forests have different understory light regimes and recruitment patterns, for example, a substantial fraction of mountain forests in New England (M211) is completely dominated by conifer species ranked as shade tolerant. Overall, mountain forests in New England and several other mountain areas across different domains, such as Mountain Provinces in the Mediterranean Division (M260) and Arizona-New Mexico Mountains Semidesert-Open Woodland–Coniferous Forest–Alpine Meadow Province (M313) - appear in the same narrow distribution and fast convergence group (Figure 8). At the same time mountain forests of Oregon and Washington (M240, Marine Division Mountain Provinces), located in the western part of the Humid Temperature Domain, display a distinctive pattern of wide distribution and slow convergence. The low number of available remeasured plots (less than 150) in northern provinces within the Dry Domain did not allow us to conclusively employ our methodology. In particular, the Temperate Steppe Division (330) as well as its mountain provinces (M330) were not classified using the spectral gap criterion (Figure 8). Forests in the southern part of the Dry Domain (Figure 8), such as Tropical Subtropical Steppe and Desert Divisions (310, 320), are classified as not being driven by shade tolerance, consistently with their low biomass, low diversity and relatively old stands completely dominated by shade intolerant species (Figure 5).

Qualitatively, the regions proposed to be shade tolerance driven exhibit also regular transition matrices, with high values concentrated primarily on the main diagonal, and secondarily on the upper diagonal above it (Figure 7). We also compared our conceptual model with the dynamics of shade tolerance index using the chronosequence approach (Figure 1 in Appendix 5). The obtained regressions complement our classification by demonstrating that ecoregions with narrow distributions do not display the expected decrease in the early stages followed by an increase during the understory reinitiation (Figure 2).

## 4 Discussion

The shade tolerance index introduced in this paper is designed as a quantitative measure of forest succession according to the classic theory based on gap dynamics and replacement of shade intolerant by shade tolerant species. This study shows that this index can be utilized to understand the forest stand dynamics in ecoregions where the classic theory is validated, as it represents forest succession scale. In particular, our results demonstrate that this index is in agreement with the continuum index developed by Curtis and McIntosh (1951) as well as the gap model simulation (Strigul et al., 2008). However, there are several challenges in the interpretation of the shade tolerance index results, which are related to existence of different successional pathways, disturbance regime complexity, spatial heterogeneity and non-stationarity of environmental variables.

The shade tolerance axis computed for southern Wisconsin data is amazingly similar to the continuum index axis developed by Curtis and McIntosh (1951). In the work cited the authors have derived the continuum index and the climax adaptation numbers based on extensive empirical observations of forest succession patterns in southern Wisconsin and measurements of species relative abundance within the stand without using the concepts of shade tolerance and gap dynamics. The shade tolerance index is based on the mechanistic succession model, as opposed to an empirically defined continuum index. Several attempts to redefine the continuum index following Curtis and McIntosh (1951) in other ecoregions have produced mixed results, unsuccessful (Buell et al., 1966) and successful (Nakamura, 1985). The major problem when developing a continuum index is the existence of several alternative successional pathways which coexist within the same mosaic of forest stands (Buell et al., 1966, Kessell and Potter, 1980). Indeed, Curtis and McIntosh (1951) considered only one successional pathway to create their continuum index and they were successful, as this was the primary successional pathway in southern Wisconsin (Buell et al., 1966). Similarly, Nakamura (1985) has successfully applied similar technique to study the only *Larix-Abies-Tsuga* successional pathway on the Mount Fuji forest. However, Kessell and Potter (1980) demonstrated the existence of two distinct successional pathways that branch from the same conditions, and obviously the continuum index cannot be constructed in this case. Overall, the existence of several successional pathways is considered as a major challenge for modeling of forest succession (Leps, 1988, Ashton, 1992, Ashton et al., 2001, Pickett et al., 1987, Johnstone and Chapin III, 2006). The advantage of the shade tolerance index is that it is based on a mechanism operating independently from the species composition of forest plots. The index can be used to describe the successional patterns in ecoregions where several alternative species replacement pathways coexist. In this case, a direct inversion of the shade tolerance index is not possible as different plots may have the same shade tolerance index level and very different species composition.

Our study of the FIA database led to the observation of highly conserved correlation patterns across time and space (Figures 6 in main text and Figures 3, 5 and 4 in Appendix 4). It is worth emphasizing that (a) the correlation structure is similar across different ecoregions, and that (b) it is not affected by the change in the forest sampling protocol that was adopted nationwide in 1999. It is further highly remarkable that the correlation structure is amazingly similar to the one obtained in Eastern Canada in a previous work (Lienard et al., 2014), despite substantial differences between the two forest inventories (e.g. climatic variables, species compositions, permanent plot designs, sampling protocols, experimental method used to determine the age of trees, stand biomass calculation). Overall, the preserved correlation patterns point toward important structural similarities between forested ecosystems in most of North America, and support the view that the shade tolerance index is a meaningful dimension that completes the usual indicators of biomass, basal area, implied stand age, and biodiversity.

We study the dynamics of the shade tolerance index in a Markov chain framework applied to the forest stand mosaic (Strigul et al., 2012, Lienard et al., 2014). Our approach is however fundamentally different from prior works using Markov chains to model succession as a change of species composition. For example, traditional Markov chain models (Usher, 1979) are based on expert knowledge of species transitions; empirically-based models (Waggoner and Stephens, 1970, Stephens and Waggoner, 1980) are deduced from a substantial collection of species observations under the assumption that the Markov chain is stationary; mechanistically-based models (Horn, 1974, 1981) predict overstory changes using a detailed survey of the understory. Markov chain models were also applied on longer time scale to predict forest type transitions (Logofet and Lesnaya, 2000, Korotkov et al., 2001). In contrast with the prior models, we consider the shade tolerance index instead of the specific species composition, as the former measure describes a forest stand in a more generic way than the latter. In particular, this approach allows the comparison of succession in ecoregions with highly different species composition.

Our analysis indicates that the classic succession theory is not universally valid for all ecoregions (Figure 8). Moreover, it reveals that shade tolerance driven succession is linked with climatic and landscape factors that determine the division level classification of ecoregions (Bailey, 1995, 2004). The most evident result concerns the eastern part of the Humid Temperate Domain, where we report a North–South monotonic transition from shade tolerance driven succession to other types of forest succession. This transition pattern is even more explicit if we exclude from consideration mountain areas, where the patch mosaic analysis should be modified to include altitude related climatic variables (Section 3.2 and Figure 2 in Appendix 4). We hypothesize that the observed transition is caused by the climatic transition to more arid conditions (Bailey, 1995). This is partially supported by the fact that shade tolerance ranking is negatively correlated with the drought tolerance ranking (Niinemets and Valladares, 2006). Another supporting fact is that southernmost forests in the Dry domain are completely dominated by shade intolerant species (see Section 3.2). Our classification method is data dependent and we were able to investigate provinces with more than 150 resampled plots available. This restriction did not allow us to make a definite statement about shade tolerance driven succession in the western part of the Humid Temperate Domain and Nothern part of the Dry Domain where mountain areas are extended. However, the classification based on only one criterion, the width of the shade tolerance distribution (Figure 8), indicates that shade tolerance may play an important role in these ecosystems. Overall, our analysis demonstrates that shade tolerance succession can be quantified at the landscape scale and connected with climatic variables. We also anticipate that this methods can be generalized and applied to other successional mechanisms.

The gap dynamics and replacement of shade intolerant species by shade tolerant species is the essential successional mechanism employed in forest gap models such as JABOWA-FORET, SPACE, ZELIG and SORTIE models, their modifications and alternatives (Shugart, 1984, Huston and Smith, 1987, Busing, 1991, Pacala et al., 1993, Pacala and Deutschman, 1995, Acevedo et al., 1995, Bugmann, 2001, Busing and Mailly, 2004, Strigul et al., 2008, Strigul, 2012). These models simulate forest succession using the growth/mortality trade-off between fast growing shade intolerant early successional species and slower growing shade tolerant species as a mechanism of tree replacement and canopy recruitment (Shugart, 1984, Kobe et al., 1995, Deutschman et al., 1999). In particular, the computer model that we used (Strigul et al., 2008) follows idealized succession dynamics that rely mostly on gap-filling mechanisms and light-driven competition. Scaling methods approximating forest gap dynamics also employs the shade tolerance driven forest dynamics as the primary successional mechanism. Kohyama (1993, 2006) has employed the patch-mosaic concept for scaling forest gap dynamics to the landscape level. In the Kohyama model the forest succession is driven by the simple birth-and-disaster process presented as the conservation equation proposed by Levin and Paine (1974). The current study may call for substantial modifications of individual based models and for their validation in ecoregions where shade tolerance is not the main mechanism of succession.

## 5 Conclusion

In this paper, we have reappraised the classic theory that shade tolerance is a main determinant in forest succession. The shade tolerance axis defined according to the classic succession paradigm demonstrates amazing similarities with other successional models such as the empirically defined continuum index which captures successional patterns in southern Wisconsin, and the successional dynamics of the two-species system of White Pine—Eastern Hemlock in the northeastern US. We further studied the spatiotemporal correlations between shade tolerance and other stand level characteristics across the US ecoregions using a data-intensive approach, and found that the shade tolerance index is uncorrelated with biomass, basal area, implied stand age, and biodiversity. The shade tolerance index allows us to test the validity of shade tolerance driven succession in particular ecoregions. Based on the distribution of shade tolerance across stands as well as their convergence rate toward equilibrium, we derived a classification of US ecoregions. Shade tolerance driven succession appears to be strictly connected with climatic variables. The importance of this successional mechanism decreases along the transition from North to South within the eastern part of the Humid Temperate Domain, and, evidently, shade tolerance is not related to successional patterns in arid ecosystems such as the southern part of the Dry Domain.

## 6 Acknowledgements

This work was partially supported by a grant from the Simons Foundation (#283770 to N.S.) and a Washington State University New Faculty SEED grant. We are grateful to Thomas Lonon and David Vaccari who kindly edited the last version of the manuscript.

## 7 Appendices

(Appendix 1) Statistical Analysis of Forest Inventory Data. Computation of the Shade Tolerance Index

(Appendix 2) Analysis of Forest Succession in southern Wisconsin

(Appendix 3) Succession in White Pine – Eastern Hemlock forests

(Appendix 4) Correlation Analysis of Forest Stand Characteristics across the US ecoregions

(Appendix 5) Supplementary Results for the classification of the US Ecoregions

(Appendix 6) Shade Tolerance Rankings for Tree Species in the FIA Database

## APPENDIX 1 Statistical Analysis of Forest Inventory Data and Computation of the Shade Tolerance Index

### 1 Statistical analyzes of the FIA database

The Forest Inventory and Analysis (FIA) is a forest survey program of the US department of Agriculture (Forest Inventory and Analysis Program, 2010). In this study employs a freely available FIA database that was downloaded from: http://www.fia.fs.fed.us/. Standard stand-level characteristics (biomass, stand age, and basal area) were computed according to the FIA manual (Forest Inventory and Analysis Program, 2010), other commonly used forest characteristics (Gini-Simpson and Shannon diversities) were computed according the procedures described elsewhere (Strigul et al., 2012, Lienard et al., 2014), and the computation of the original shade tolerance index is described in the next section. In particular, we extracted the following information about the trees of each plot, from Table `TREE`:

- The status of each tree (`STATUSCD`). We analyzed only trees with this variable set to 1 as we consider only live trees in this study.
- The carbon estimate above ground (`CARBON_AG`) and below ground (`CARBON_BG`). These variables are used to deduce the biomass as: 2 *×* (`CARBON_AG + CARBON_BG`).
- The diameter at breast height (`DIA`), used to compute the basal area as: *π×* `DIA` ^2^.
- The tree species (`SPCD`), used to link with the shade tolerance ranking table, provided as a supplemental file.
- For the analysis of plot dynamics, we used the year of the survey (`INVYR`), as well as locations variables used in the computation of a unique identifier for each plot (`PLOT, COUNTYCD, UNITCD, STATECD`).
- The scaling factor (`TPA_UNADJ`), allowing to normalize plots of different sizes.

We also extracted information of plots from Table `COND`:

- The stand age (`STDAGE`), which is computed as the average of trees in a stand. We refer to this variable as the implied stand age in the text of the manuscript.
- The soil type (`PHYSCLCD`), used to select plots located on mesic soils.

In addition to these variables, we extracted from Table `PLOT` the following:

- Latitude (`LAT`) and longitude (`LON`) of plots.
- The ecoregion code (`ECOSUBCD`), from which we retained only the first four characters to derive Bailey’s provinces (including a possible starting space, e.g., “M211” or “211”). See Figure 1 in Appendix 4 for a map of the provinces as reported in the database.

The Gini-Simpson diversity index (Simpson, 1949) is equal to 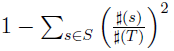, with *S* being the set of species in a stand, *♯*(*s*) being the number of trees with species s and *♯*(*T*) being the total number of trees. The Shannon diversity index (Shannon and Weaver, 1949) is equal to is equal to 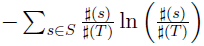, with the same notations. Both the Gini-Simpson and the Shannon indexes are in the 0-1 range, where high values indicate a high species heterogeneity.

### 2 Computation of the shade tolerance index using FIA data

We have explored several alternatives to calculate Ω*_j_*, the abundance of tree species for a stand: 1) number of trees (canopy or understory) of the species *j*, 2) sum of the basal areas of the species *j* trees (total, canopy or understory), 3) biomass of the species *j*, 4) area of canopy occupied by species *j*. However, we discovered that these estimates are highly correlated (Appendix 4). Therefore is sufficient to compute only one of alternative Ω*_j_*. In this study, we employed the abundance based on the sum of basal areas for the following reasons: a) the sum of basal areas is widely used in forestry for this purpose, b) statistically basal area is often related to crown area, and, therefore, the sum of basal areas can be also related to the canopy area occupied by species *j* (which is not recorded in the FIA dataset), c) this parameter is easy to calculate from the FIA dataset.

All the tree species in the FIA database were classified according to the shade tolerance tables (Baker, 1949, Burns and Honkala, 1990, Humbert et al., 2007) in 5 categories with respect to the shade tolerance: very intolerant, intolerant, intermediate, tolerant, and very tolerant. We assume that the scale of shade tolerance is linear and the classification of shade tolerance uniformly partition this scale, similarly to Humbert et al. (2007), Kunstler et al. (2009), Valladares and Niinemets (2008). Then we assign the following numbers to the shade tolerance classes: very intolerant = 0, intolerant = 0.25, intermediate = 0.5, tolerant = 0.75, and very tolerant = 1. The shade tolerance rankings for North American tree species along with the species code from the FIA database is provided in a comma-separated file (online Appendix 6).

### 3 Statistical description of US forests across ecoregions

**Table 1:** Shade tolerance index distribution across US provinces.

| Province | Min. | 1st Qu. | Median | Mean | 3rd Qu. | Max. |
| --- | --- | --- | --- | --- | --- | --- |
| 211 | 0 | 0.525 | 0.6794 | 0.6615 | 0.8273 | 1 |
| 212 | 0 | 0.261 | 0.4882 | 0.491 | 0.7258 | 1 |
| 221 | 0 | 0.3913 | 0.4982 | 0.4985 | 0.6006 | 1 |
| 222 | 0 | 0.3458 | 0.4863 | 0.4691 | 0.5908 | 1 |
| 223 | 0 | 0.3316 | 0.4349 | 0.4311 | 0.5196 | 1 |
| 231 | 0 | 0.2552 | 0.327 | 0.3496 | 0.4282 | 1 |
| 232 | 0 | 0.2795 | 0.3887 | 0.3971 | 0.5 | 1 |
| 234 | 0 | 0.3203 | 0.4393 | 0.4297 | 0.527 | 1 |
| 242 | 0 | 0.4829 | 0.5138 | 0.5534 | 0.6491 | 1 |
| 251 | 0 | 0.3498 | 0.4603 | 0.4538 | 0.5378 | 1 |
| 255 | 0 | 0.25 | 0.3306 | 0.3347 | 0.4569 | 1 |
| 261 | 0.07565 | 0.5 | 0.5876 | 0.6367 | 0.8038 | 1 |
| 262 | 0.119 | 0.1331 | 0.25 | 0.3391 | 0.5 | 0.6936 |
| 263 | 0.09627 | 0.5993 | 0.7001 | 0.7028 | 0.8333 | 1 |
| 313 | 0 | 0.007809 | 0.0791 | 0.1314 | 0.2243 | 0.75 |
| 315 | 0 | 0 | 0 | 0.1114 | 0.1719 | 1 |
| 321 | 0 | 0 | 0 | 0.07703 | 0.01074 | 1 |
| 322 | 0 | 0 | 0 | 0.04668 | 0.02958 | 0.7153 |
| 331 | 0 | 0.06614 | 0.25 | 0.2444 | 0.2776 | 1 |
| 332 | 0 | 0.1287 | 0.378 | 0.3534 | 0.5424 | 1 |
| 341 | 0 | 0 | 0.008568 | 0.09427 | 0.118 | 1 |
| 342 | 0 | 0.003473 | 0.25 | 0.2192 | 0.25 | 1 |
| 411 | 0 | 0.5 | 0.5 | 0.5331 | 0.5958 | 0.75 |
| M211 | 0 | 0.6618 | 0.7921 | 0.7592 | 0.8878 | 1 |
| M221 | 0 | 0.3925 | 0.4833 | 0.4824 | 0.562 | 1 |
| M223 | 0 | 0.2758 | 0.3915 | 0.3745 | 0.4763 | 0.8987 |
| M231 | 0 | 0.1785 | 0.2636 | 0.2782 | 0.3617 | 1 |
| M242 | 0 | 0.4515 | 0.5483 | 0.5623 | 0.7344 | 1 |
| M261 | 0 | 0.2728 | 0.4666 | 0.445 | 0.5852 | 1 |
| M262 | 0 | 0.146 | 0.2983 | 0.3248 | 0.5 | 0.75 |
| M313 | 0 | 0.043 | 0.2029 | 0.1986 | 0.2825 | 0.9106 |
| M331 | 0 | 0.1087 | 0.3447 | 0.355 | 0.5503 | 1 |
| M332 | 0 | 0.2504 | 0.4414 | 0.405 | 0.5223 | 0.9818 |
| M333 | 0 | 0.3139 | 0.4959 | 0.4826 | 0.6834 | 1 |
| M334 | 0 | 0.25 | 0.25 | 0.2625 | 0.25 | 0.6526 |
| M341 | 0 | 0 | 0.1007 | 0.1831 | 0.313 | 1 |

**Table 2:** Summary statistics of Biomass (in 10^3^kg/ha) for each US province.

| Province | Min. | 1st Qu. | Median | Mean | 3rd Qu. | Max. |
| --- | --- | --- | --- | --- | --- | --- |
| 211 | 0.09628 | 60.68 | 111.2 | 119.3 | 166.6 | 528.7 |
| 212 | 0.03543 | 41.21 | 76.53 | 84.58 | 119 | 542.5 |
| 221 | 0.08031 | 73.54 | 125.9 | 131.1 | 178.7 | 526.6 |
| 222 | 0.05631 | 39.66 | 80.04 | 90.39 | 127.4 | 521.1 |
| 223 | 0.08261 | 58.56 | 96.3 | 101.7 | 135.9 | 566 |
| 231 | 0.04592 | 49.93 | 97.01 | 105.9 | 149.7 | 557.5 |
| 232 | 0.07411 | 35.8 | 80.13 | 97.69 | 141.2 | 808.9 |
| 234 | 0.1585 | 45.57 | 97.58 | 116 | 165.8 | 870.3 |
| 242 | 0.08631 | 54.75 | 144.1 | 192.2 | 279.4 | 973.7 |
| 251 | 0.05393 | 39.06 | 73.08 | 81.18 | 114.1 | 578.3 |
| 255 | 0.1218 | 17.97 | 43.14 | 54.51 | 79.36 | 308.8 |
| 261 | 0.2588 | 40.05 | 125.7 | 216.6 | 324.5 | 1271 |
| 262 | 11 | 27.37 | 80.69 | 84.18 | 144.3 | 157.5 |
| 263 | 1.024 | 125.2 | 227.4 | 275.1 | 360.1 | 1787 |
| 313 | 0.1743 | 11.6 | 24.82 | 37.13 | 46.89 | 464.2 |
| 315 | 0.08919 | 5.123 | 11.46 | 18.71 | 25.05 | 191.5 |
| 321 | 0.08249 | 2.154 | 5.369 | 11.19 | 11.96 | 360.8 |
| 322 | 0.1633 | 6.199 | 13.32 | 20.59 | 25.14 | 133.5 |
| 331 | 0.04699 | 10.98 | 25.6 | 37.13 | 50.39 | 302.1 |
| 332 | 0.1892 | 24.7 | 48.04 | 61.57 | 83.42 | 608.1 |
| 341 | 0.08775 | 12.17 | 24.78 | 32.78 | 42.84 | 337.5 |
| 342 | 0.08842 | 7.497 | 16.55 | 30.76 | 37.79 | 377.9 |
| 411 | 0.1656 | 21.61 | 56.92 | 72.16 | 102.9 | 312.6 |
| M211 | 0.04047 | 72.11 | 121.2 | 125.3 | 171.2 | 457.7 |
| M221 | 0.08035 | 94.63 | 141.8 | 146.8 | 192.5 | 588.2 |
| M223 | 0.5212 | 70.45 | 107.9 | 108.9 | 146.9 | 378.1 |
| M231 | 0.1051 | 47.92 | 84.5 | 90.49 | 125.8 | 329.1 |
| M242 | 0.09882 | 71.22 | 178.2 | 247.1 | 363.4 | 1651 |
| M261 | 0.07831 | 49.36 | 125.9 | 174.1 | 250.2 | 1287 |
| M262 | 0.205 | 16.16 | 44.43 | 65.55 | 98.48 | 355.7 |
| M313 | 0.1894 | 14.17 | 34.04 | 51.58 | 69.75 | 368.6 |
| M331 | 0.1049 | 29.12 | 62.85 | 80.99 | 115.3 | 500.3 |
| M332 | 0.08631 | 37.21 | 78.85 | 96.64 | 133.5 | 650.1 |
| M333 | 0.1137 | 45.97 | 99.03 | 121.6 | 168.8 | 670.7 |
| M334 | 0.1024 | 31.52 | 57.61 | 66.83 | 91.94 | 275.3 |
| M341 | 0.1848 | 16.04 | 30.98 | 41.49 | 54.03 | 413.3 |

**Table 3:** Summary statistics of Basal area (in m^2^/ha) for each US province.

| Province | Min. | 1st Qu. | Median | Mean | 3rd Qu. | Max. |
| --- | --- | --- | --- | --- | --- | --- |
| 211 | 0.07509 | 14.47 | 22.96 | 22.67 | 30.36 | 75.17 |
| 212 | 0.03755 | 11.75 | 19.37 | 19.84 | 26.98 | 101.9 |
| 221 | 0.07509 | 13.19 | 20.45 | 20.28 | 26.73 | 66.74 |
| 222 | 0.03755 | 9.036 | 16.36 | 16.87 | 23.57 | 67.26 |
| 223 | 0.04544 | 12.09 | 18.19 | 17.88 | 23.46 | 71.03 |
| 231 | 0.07504 | 11.93 | 19.74 | 19.52 | 26.54 | 80.53 |
| 232 | 0.07488 | 9.098 | 17.36 | 18.87 | 26.42 | 141 |
| 234 | 0.09386 | 10.15 | 18.39 | 19.51 | 26.71 | 130.7 |
| 242 | 0.09386 | 11.38 | 23.71 | 26.94 | 38.09 | 105.8 |
| 251 | 0.03755 | 8.609 | 15.01 | 15.53 | 21.43 | 86.31 |
| 255 | 0.09386 | 5.324 | 11.27 | 12.35 | 18.02 | 57.32 |
| 261 | 0.1136 | 9.35 | 20.13 | 31.18 | 45.23 | 138.3 |
| 262 | 1.089 | 5.561 | 9.014 | 16.4 | 30.75 | 35.58 |
| 263 | 0.5876 | 23.87 | 39.8 | 42.59 | 56.86 | 154.5 |
| 313 | 0.09386 | 8.185 | 15.13 | 17.64 | 24.69 | 79.7 |
| 315 | 0.09386 | 2.876 | 5.97 | 8.238 | 11.52 | 59.87 |
| 321 | 0.09386 | 1.736 | 3.961 | 6.371 | 8.279 | 70.69 |
| 322 | 0.3184 | 4.724 | 9.115 | 12.54 | 16.43 | 70.77 |
| 331 | 0.09386 | 5.157 | 10.33 | 12.54 | 17.77 | 90.88 |
| 332 | 0.1089 | 6.418 | 12.36 | 14.09 | 19.75 | 95.97 |
| 341 | 0.09386 | 9.174 | 16.84 | 19.16 | 26.65 | 83.45 |
| 342 | 0.09386 | 4.411 | 8.852 | 12.41 | 16.7 | 98.66 |
| 411 | 0.1884 | 6.892 | 16.27 | 19.38 | 27.69 | 66.15 |
| M211 | 0.07509 | 17.11 | 24.83 | 24.47 | 31.81 | 64.18 |
| M221 | 0.07513 | 17.46 | 23.66 | 23.47 | 29.6 | 68.54 |
| M223 | 0.2713 | 14.66 | 19.22 | 19.12 | 24.36 | 47.45 |
| M231 | 0.09386 | 12.12 | 18.24 | 18.24 | 24.3 | 49.02 |
| M242 | 0.1136 | 16.96 | 31.7 | 35.26 | 49.75 | 139.1 |
| M261 | 0.1136 | 12.66 | 25.88 | 29.55 | 42.74 | 136.4 |
| M262 | 0.3692 | 5.42 | 10.56 | 14.76 | 21.74 | 62.99 |
| M313 | 0.2112 | 9.843 | 17.68 | 19.05 | 26.07 | 76.69 |
| M331 | 0.1136 | 11.54 | 20.18 | 22.27 | 31.17 | 104.8 |
| M332 | 0.09386 | 10.95 | 19.48 | 21.23 | 29.29 | 108.1 |
| M333 | 0.1136 | 11.87 | 21.73 | 23.71 | 32.54 | 106.6 |
| M334 | 0.2117 | 9.553 | 16.11 | 16.83 | 22.45 | 49.31 |
| M341 | 0.1136 | 10.21 | 18.69 | 20.6 | 28.45 | 92.67 |

**Table 4:** Summary statistics of Average age of trees (implied stand age) for each US province.

| Province | Min. | 1st Qu. | Median | Mean | 3rd Qu. | Max. |
| --- | --- | --- | --- | --- | --- | --- |
| 211 | 1 | 45 | 65 | 62.62 | 82 | 195 |
| 212 | 1 | 35 | 55 | 54.99 | 73 | 272 |
| 221 | 1 | 43 | 59 | 57.54 | 73 | 203 |
| 222 | 1 | 35 | 51 | 54.18 | 70 | 311 |
| 223 | 1 | 40 | 56 | 56.16 | 71 | 162 |
| 231 | 1 | 16 | 35 | 37.65 | 55 | 195 |
| 232 | 1 | 15 | 33 | 38.87 | 55 | 195 |
| 234 | 1 | 30 | 50 | 46.92 | 64 | 170 |
| 242 | 1 | 21 | 45 | 49.64 | 70 | 350 |
| 251 | 1 | 35 | 50 | 52.35 | 68 | 206 |
| 255 | 1 | 25 | 40 | 41.85 | 55 | 130 |
| 261 | 2 | 74.75 | 90 | 113.4 | 126.2 | 600 |
| 262 | 50 | 60 | 60 | 70 | 80 | 100 |
| 263 | 1 | 40 | 55 | 67.6 | 80 | 500 |
| 313 | 2 | 93 | 134 | 141.9 | 184.5 | 522 |
| 315 | 1 | 25 | 40 | 45.04 | 55 | 550 |
| 321 | 1 | 25 | 45 | 66.68 | 92 | 318 |
| 322 | 5 | 77 | 110 | 121.9 | 160 | 390 |
| 331 | 1 | 50 | 74 | 75.7 | 95 | 328 |
| 332 | 3 | 35 | 50 | 50.94 | 65 | 130 |
| 341 | 2 | 90 | 130.5 | 137.9 | 180 | 1028 |
| 342 | 4 | 61 | 83 | 98.1 | 120 | 380 |
| 411 | 1 | 30 | 50 | 55.46 | 82 | 139 |
| M211 | 1 | 46 | 66 | 65.17 | 83 | 238 |
| M221 | 1 | 45 | 64 | 61.91 | 80 | 200 |
| M223 | 2 | 50 | 63 | 58.54 | 71 | 111 |
| M231 | 1 | 27 | 49 | 46.32 | 65 | 115 |
| M242 | 1 | 31 | 65 | 96.16 | 120 | 800 |
| M261 | 1 | 62 | 89 | 106.3 | 125 | 997 |
| M262 | 2 | 70 | 94 | 110.8 | 150 | 325 |
| M313 | 1 | 80 | 103 | 109.7 | 134 | 343 |
| M331 | 1 | 67 | 98 | 102.2 | 134 | 733 |
| M332 | 2 | 70 | 98 | 104.8 | 133 | 494 |
| M333 | 1 | 48 | 79 | 82.87 | 109 | 404 |
| M334 | 1 | 67 | 85 | 84.21 | 100 | 278 |
| M341 | 2 | 78.75 | 118.5 | 125.7 | 168 | 548 |

**Table 5:** Summary statistics of Shannon diversity index for each US province.

| Province | Min. | 1st Qu. | Median | Mean | 3rd Qu. | Max. |
| --- | --- | --- | --- | --- | --- | --- |
| 211 | 0 | 0.8676 | 1.197 | 1.148 | 1.476 | 2.242 |
| 212 | 0 | 0.6315 | 0.9831 | 0.9409 | 1.297 | 2.3 |
| 221 | 0 | 1.029 | 1.38 | 1.311 | 1.666 | 2.571 |
| 222 | 0 | 0.6909 | 1.089 | 1.043 | 1.43 | 2.465 |
| 223 | 0 | 1.169 | 1.512 | 1.44 | 1.779 | 2.694 |
| 231 | 0 | 0.9968 | 1.411 | 1.315 | 1.725 | 2.689 |
| 232 | 0 | 0.5228 | 0.9988 | 0.9405 | 1.395 | 2.509 |
| 234 | 0 | 0.7053 | 1.176 | 1.094 | 1.508 | 2.35 |
| 242 | 0 | 0.2206 | 0.6525 | 0.6472 | 1.03 | 1.709 |
| 251 | 0 | 0.76 | 1.192 | 1.134 | 1.542 | 2.657 |
| 255 | 0 | 0.4698 | 0.8777 | 0.8464 | 1.267 | 2.435 |
| 261 | 0 | 0 | 0.5943 | 0.5254 | 0.868 | 1.436 |
| 262 | 0 | 0 | 0.153 | 0.4195 | 0.8897 | 1.055 |
| 263 | 0 | 0.5656 | 0.8039 | 0.7863 | 1.048 | 1.648 |
| 313 | 0 | 0.1011 | 0.5236 | 0.4721 | 0.687 | 1.672 |
| 315 | 0 | 0 | 0 | 0.2756 | 0.5727 | 1.607 |
| 321 | 0 | 0 | 0 | 0.2322 | 0.4978 | 1.555 |
| 322 | 0 | 0 | 0.216 | 0.303 | 0.5985 | 1.446 |
| 331 | 0 | 0 | 0.1838 | 0.3134 | 0.6277 | 1.604 |
| 332 | 0 | 0.3386 | 0.6913 | 0.6951 | 1.053 | 1.822 |
| 341 | 0 | 0 | 0.4325 | 0.3832 | 0.6497 | 1.557 |
| 342 | 0 | 0 | 0 | 0.156 | 0.1753 | 1.375 |
| 411 | 0 | 0 | 0.4862 | 0.512 | 0.8577 | 1.891 |
| M211 | 0 | 0.9022 | 1.214 | 1.175 | 1.49 | 2.272 |
| M221 | 0 | 1.168 | 1.471 | 1.412 | 1.719 | 2.477 |
| M223 | 0 | 1.251 | 1.547 | 1.486 | 1.77 | 2.421 |
| M231 | 0 | 0.9651 | 1.338 | 1.263 | 1.621 | 2.356 |
| M242 | 0 | 0.3185 | 0.6653 | 0.6375 | 0.9483 | 1.871 |
| M261 | 0 | 0.2479 | 0.6667 | 0.636 | 0.9777 | 1.858 |
| M262 | 0 | 0 | 0.3073 | 0.3629 | 0.6277 | 1.327 |
| M313 | 0 | 0.2105 | 0.6365 | 0.5933 | 0.911 | 1.766 |
| M331 | 0 | 0.08903 | 0.5353 | 0.4963 | 0.7655 | 1.714 |
| M332 | 0 | 0.1124 | 0.5323 | 0.5088 | 0.7968 | 1.7 |
| M333 | 0 | 0.3652 | 0.6894 | 0.691 | 1.015 | 1.871 |
| M334 | 0 | 0 | 0 | 0.2283 | 0.4491 | 1.358 |
| M341 | 0 | 0.1884 | 0.5598 | 0.4994 | 0.6906 | 1.596 |

**Table 6:** Summary statistics of Gini-Simpson diversity index for each US province.

| Province | Min. | 1st Qu. | Median | Mean | 3rd Qu. | Max. |
| --- | --- | --- | --- | --- | --- | --- |
| 211 | 0 | 0.475 | 0.6209 | 0.5697 | 0.7181 | 0.8788 |
| 212 | 0 | 0.3529 | 0.5423 | 0.4893 | 0.668 | 0.8902 |
| 221 | 0 | 0.5384 | 0.6753 | 0.6208 | 0.762 | 0.9083 |
| 222 | 0 | 0.4069 | 0.5892 | 0.527 | 0.7068 | 0.9011 |
| 223 | 0 | 0.6036 | 0.72 | 0.6665 | 0.7901 | 0.9227 |
| 231 | 0 | 0.538 | 0.6941 | 0.6227 | 0.7799 | 0.9256 |
| 232 | 0 | 0.2981 | 0.5558 | 0.4815 | 0.7014 | 0.9075 |
| 234 | 0 | 0.415 | 0.6102 | 0.537 | 0.7252 | 0.8926 |
| 242 | 0 | 0.1072 | 0.4087 | 0.3635 | 0.5917 | 0.7844 |
| 251 | 0 | 0.4451 | 0.625 | 0.5588 | 0.7349 | 0.9202 |
| 255 | 0 | 0.2482 | 0.5 | 0.4466 | 0.6616 | 0.8967 |
| 261 | 0 | 0 | 0.3455 | 0.305 | 0.4999 | 0.7388 |
| 262 | 0 | 0 | 0.06827 | 0.2449 | 0.5161 | 0.64 |
| 263 | 0 | 0.3121 | 0.4952 | 0.4425 | 0.5998 | 0.791 |
| 313 | 0 | 0.03957 | 0.32 | 0.2908 | 0.48 | 0.7929 |
| 315 | 0 | 0 | 0 | 0.1671 | 0.3627 | 0.7812 |
| 321 | 0 | 0 | 0 | 0.1385 | 0.3013 | 0.7615 |
| 322 | 0 | 0 | 0.1049 | 0.1915 | 0.3912 | 0.7435 |
| 331 | 0 | 0 | 0.08318 | 0.1924 | 0.4086 | 0.7978 |
| 332 | 0 | 0.1725 | 0.4444 | 0.3872 | 0.595 | 0.8281 |
| 341 | 0 | 0 | 0.2512 | 0.2459 | 0.4444 | 0.78 |
| 342 | 0 | 0 | 0 | 0.09683 | 0.0811 | 0.7445 |
| 411 | 0 | 0 | 0.2864 | 0.2928 | 0.5089 | 0.8195 |
| M211 | 0 | 0.4918 | 0.6289 | 0.5844 | 0.723 | 0.882 |
| M221 | 0 | 0.5958 | 0.707 | 0.661 | 0.7768 | 0.905 |
| M223 | 0 | 0.6358 | 0.7324 | 0.6912 | 0.7902 | 0.9031 |
| M231 | 0 | 0.5297 | 0.6779 | 0.6156 | 0.758 | 0.8993 |
| M242 | 0 | 0.1599 | 0.4182 | 0.3648 | 0.5468 | 0.8228 |
| M261 | 0 | 0.1171 | 0.4061 | 0.3559 | 0.5582 | 0.8155 |
| M262 | 0 | 0 | 0.1618 | 0.216 | 0.4102 | 0.6798 |
| M313 | 0 | 0.0986 | 0.3969 | 0.3434 | 0.5301 | 0.8178 |
| M331 | 0 | 0.0334 | 0.3171 | 0.2938 | 0.493 | 0.7902 |
| M332 | 0 | 0.04519 | 0.3134 | 0.2981 | 0.4974 | 0.8047 |
| M333 | 0 | 0.19 | 0.4322 | 0.3839 | 0.5714 | 0.8264 |
| M334 | 0 | 0 | 0 | 0.1386 | 0.2604 | 0.7033 |
| M341 | 0 | 0.08588 | 0.3476 | 0.3074 | 0.4851 | 0.7901 |

### 4 Spatial visualization of forest characteristics in decades

**Figure 1:**
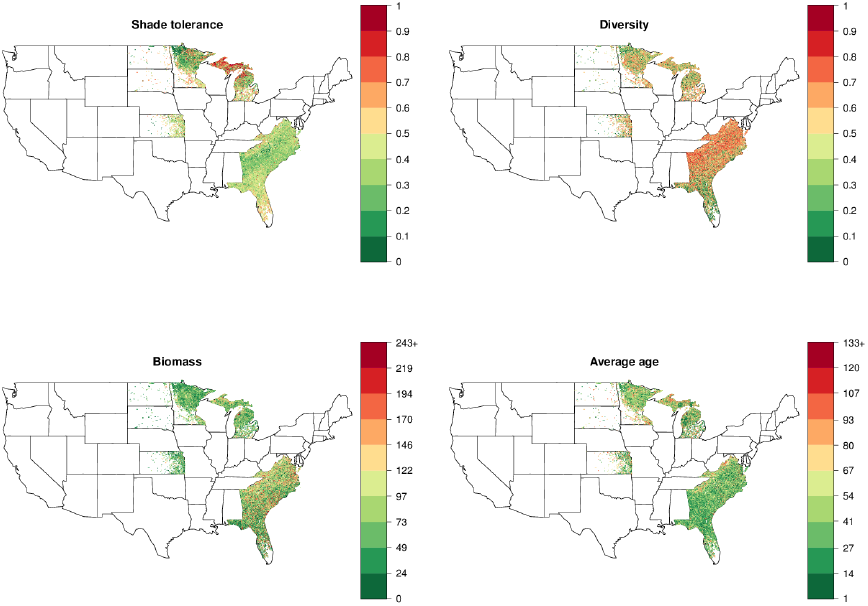
Stand-level characteristics of plots for 1968-1982.

**Figure 2:**
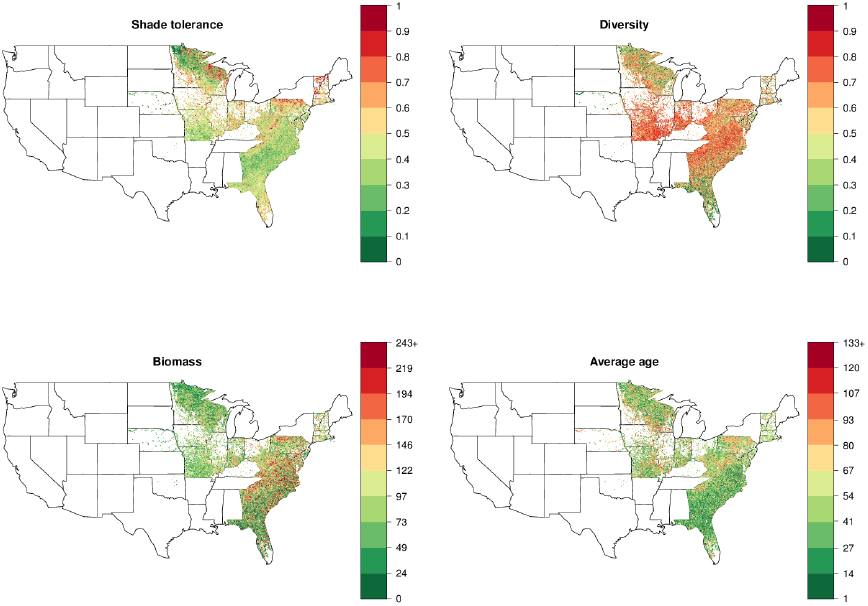
Stand-level characteristics of plots for 1982-1992.

**Figure 3:**
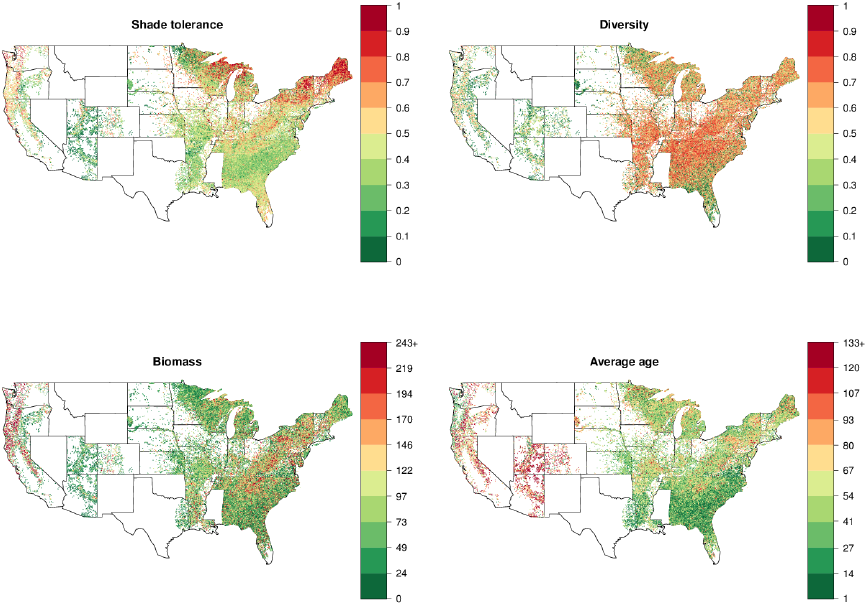
Stand-level characteristics of plots for 1992-2002.

**Figure 4:**
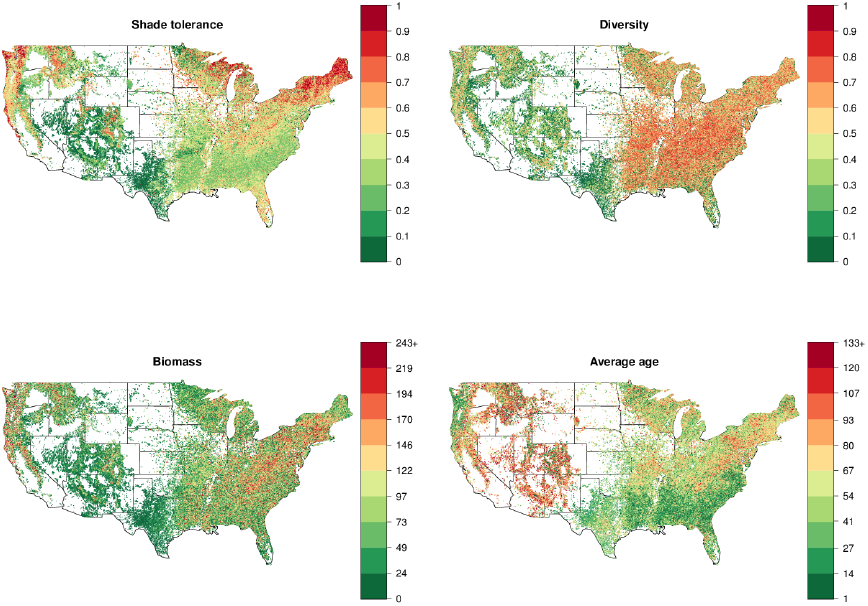
Stand-level characteristics of plots for 2002-2012.

## APPENDIX 2 Analysis of Forest Succession in southern Wisconsin

**Figure 1:**
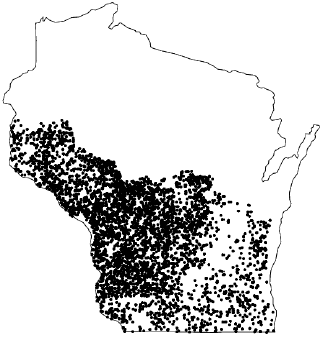
Plot locations in southern Wisconsin used in Section 2.3 for the comparison with Curtis and McIntosh (1951). The overall geographical area considered is similar to the Figure 1 in Curtis and McIntosh (1951), while the number of plots (n=7017) considered here is substantially larger than in the original study (n=95).

**Figure 2:**
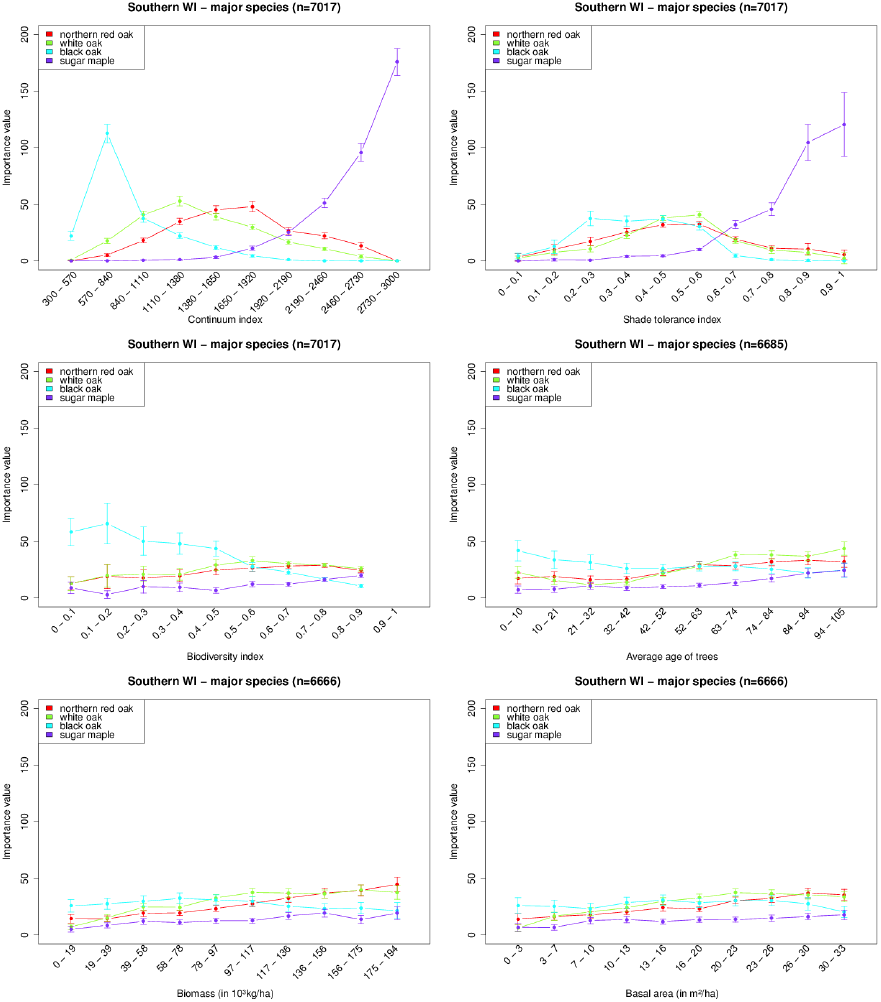
Characteristics for the major species in southern Wisconsin (Section 2.3). Bars indicate the standard error of the mean.

**Figure 3:**
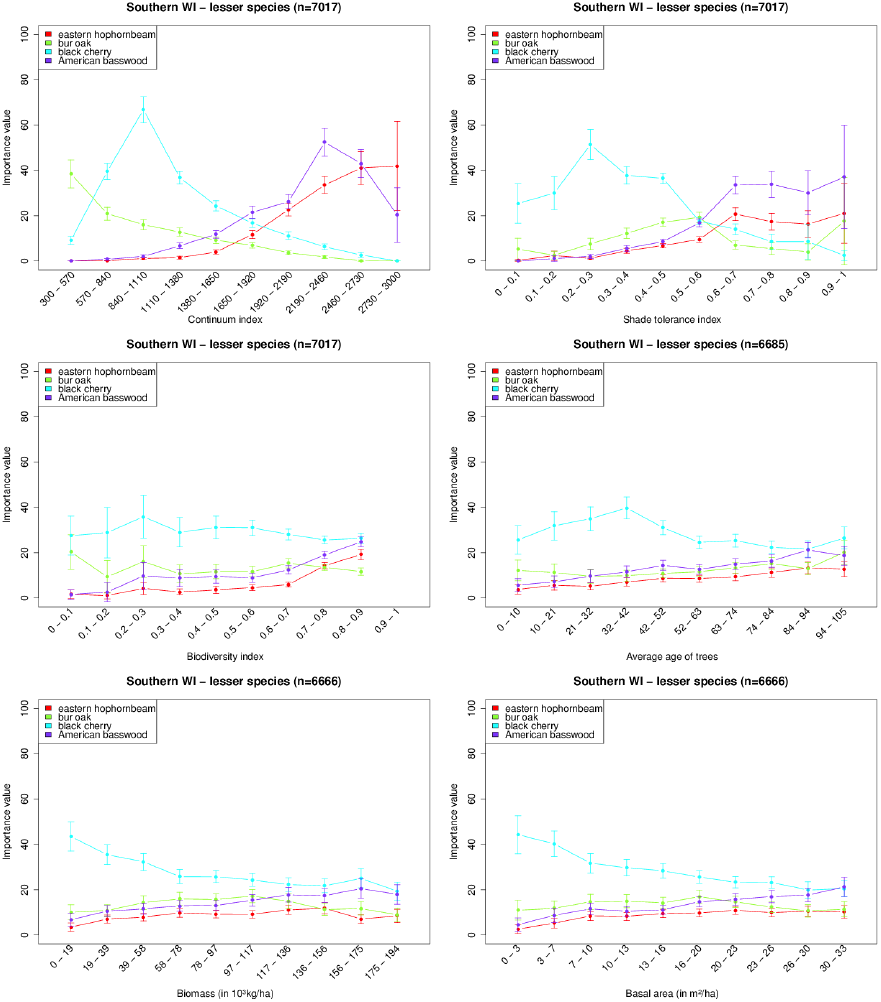
Characteristics for the lesser species in southern Wisconsin (Section 2.3). Bars indicate the standard error of the mean.

## APPENDIX 3 Succession in White Pine – Eastern Hemlock forests

**Figure 1:**
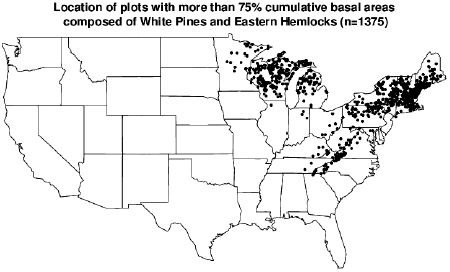
Map of the US showing the location of White Pine – Eastern Hemlock two-species systems (studied in the comparison with the computer model at Section 2.4).

**Figure 2:**
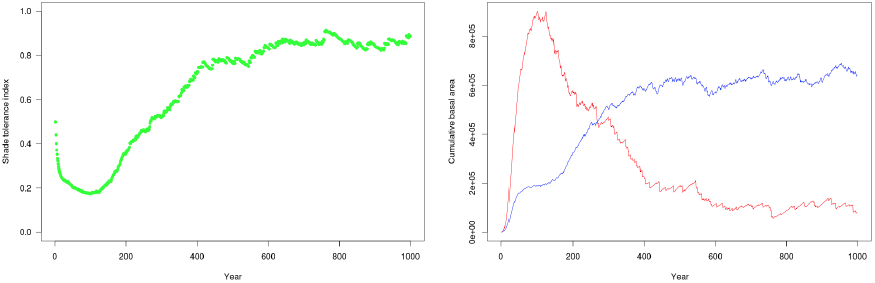
Computer simulation of one hectare containing White Pine – Eastern Hemlock forests (1 hectare, 1000 years), as in Figure 3 in main text, but with the x-axis representing time after a major disturbance. Left: shade tolerance index. Right: cumulative basal area of white pines (blue) and eastern hemlock (red).

## APPENDIX 4 Correlation Analysis of Forest Stand Characteristics across the US ecoregions

### 1 Bailey’s provincial subdivision of the US

**Figure 1:**
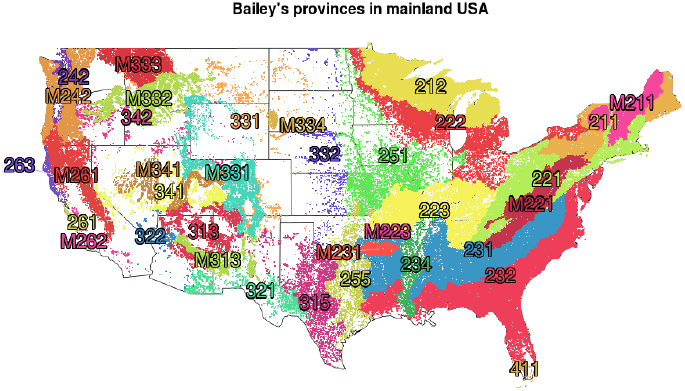
Bailey’s provinces (Bailey, 1995), as reported in the FIA database (first four characters from column `ECOSUBCD` of table `PLOT`). Each dot corresponds to one measured stand; different colors were randomly chosen to allow the distinction between provinces. The number of plots per ecoregion can be found in Table 1.

**Figure 2:**
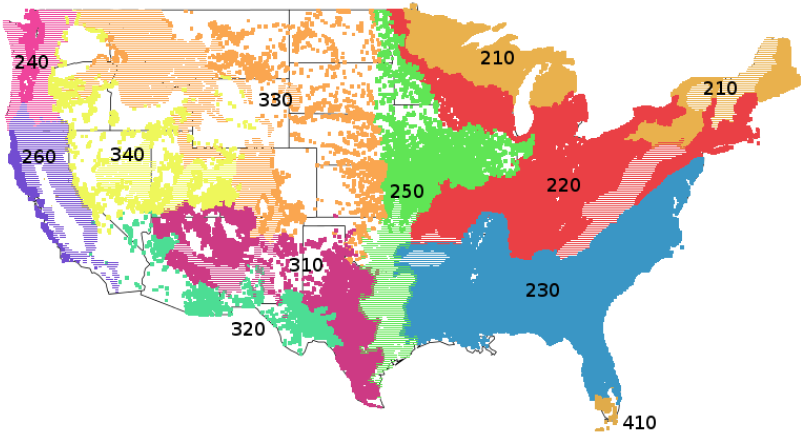
Ecoregion divisions of the US forested ecosystems (see also Bailey, 1995, and Figure 1 in this appendix). The first digit “2” indicates divisions inside the Humid Temperate Domain (e.g. 210), “3” inside the Dry Domain and “4” inside the Humid Tropical Domain. Dashed areas correspond to mountain provinces.

### 2 Correlations of different forest characteristics depending on computation methods

**Figure 3:**
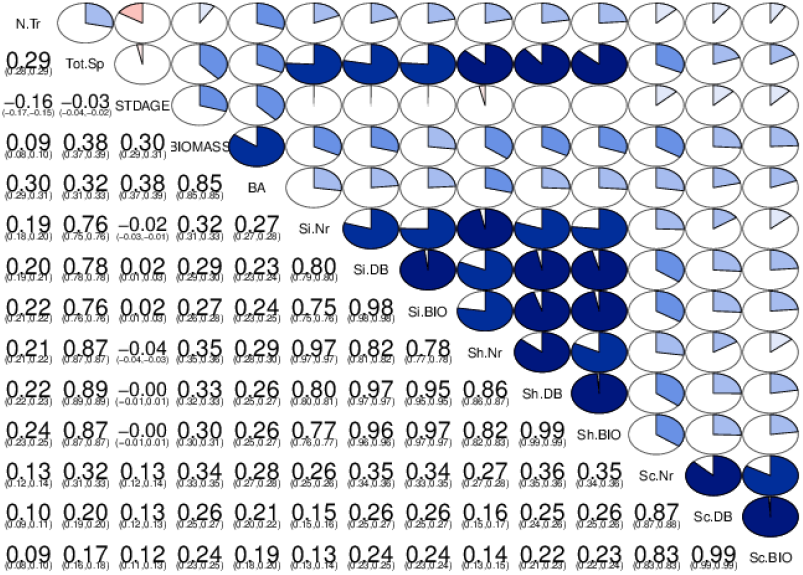
Correlations in the whole dataset using Pearson’s coefficients represented graphically (upper diagonal) and numerically with 95% confidence interval (lower diagonal) (Friendly, 2002). Abbreviations: `N.Tr` is the number of trees per hectare; `Tot.Sp` is the species richness; `STDAGE` is the stand age as reported in the FIA database; `BIOMASS` is the total biomass per hectare; `BA` is the Basal Area per hectare; `Si.Nr, Si.DB` and `Si.BIO` are the Gini-Simpson diversity indexes based on number of trees, basal area and biomass respectively; similarly, `Sh.Nr, Sh.DB` and `Sh.BIO` are Shannon diversity indexes, and `Sc.Nr, Sc.DB` and `Sc.BIO` are Shade tolerance indexes. The shade tolerance indices calculated with different species abundance measures (number of canopy trees, biomass, and basal area) are highly correlated, with a Pearson’s coefficient r > 0.8, and therefore only one of these variables can be employed.

### 3 Temporal analysis of the correlations

**Figure 4:**
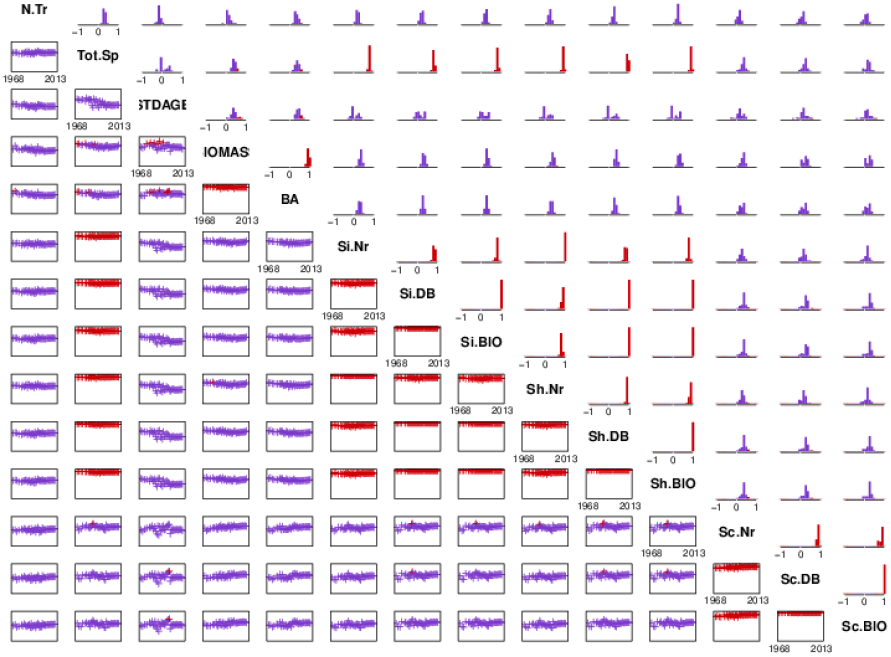
Correlations in the whole dataset broken down by years. The lower panel represents the yearly correlation coefficients as a function of time, with y-axis ranging from -1 to 1; the upper diagonal are histogram of the correlations represented in the lower diagonal. Years 1975, 1976 and 2013 were excluded as they contained less than 500 observations. Abbreviations are as in Figure 3.

### 4 Spatial analysis of the correlations

**Figure 5:**
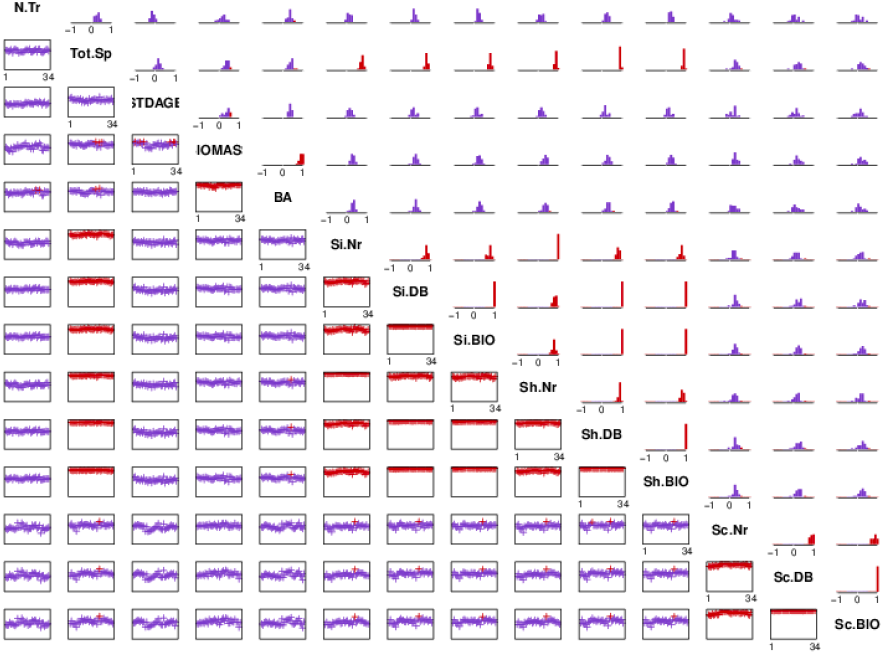
Correlations in the whole dataset broken down by ecoregion, similarly to Figure 4. Provinces 262 and 263 were excluded because they contained less than 500 observations. Abbreviations are as in Figure 3.

## APPENDIX 5 Supplementary Results for the classification of the US Ecoregions

### 1 Methodology to estimate the transition matrices

It is challenging to estimate the transition probabilities of shade tolerance in a Markov chain model based on forest inventory data, because the sampling interval of plots is highly irregular. To overcome this difficulty, we employed here the original methodology developed in Lienard et al. (2014). Specifically, we applied Gibbs sampling, a Monte Carlo Markov Chain implementation (Robert and Casella, 2004) to estimate the transition probabilities, with the specific guidelines provided by Pasanisi et al. (2012). We provide in the following a brief description of the algorithm; please refer to Lienard et al. (2014) for an extended description.

First, we constructed a temporal sequence *S_p_* of the shade tolerance index for each plot *p*, by inserting the discretized value of shade tolerance index *s*_(*p,i*)_ measured in the *i*-th year, at position *i* of *S_p_*. Each temporal sequence *S_p_* is mostly composed of unknown values, as only a fraction of the forest plots were surveyed each year. We then reduced the sparseness of these sequences by averaging the values in 3-year bins (a valid dimensionality reduction given that only % of the database recordings were spaced of less than 3 years). We further extracted sub-sequences of length 5 (thus spanning 15 years) that fulfilled three criteria: (a) each subsequence starts with a known value, (b) each subsequence contains at least two known values. Let *Y* be the matrix constructed using all the sub-sequences, with rows corresponding to successive measures of different plots and columns corresponding to different time steps. The initialization of Gibbs sampling consists of replacing the missing values in *Y* at random, resulting in so-called augmented data *Z*^[0]^. Then, the two following steps are iterated *H* = 500 times:

1. in the **parameter estimation** step, we draw a new transition matrix 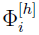 conditional on the augmented data *Z^h−^*^1^:

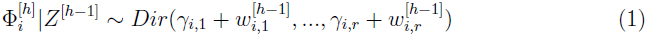 with *Dir* is the Dirichlet distribution, *γ* are biasing factors set here uniformly to 1 as we include no prior knowledge on the shape of the transition matrix (Pasanisi et al., 2012), and *w_i,j_* are the sufficient statistics defined as

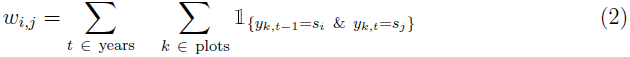
2. in the **data augmentation** step, we draw new values for the missing states:

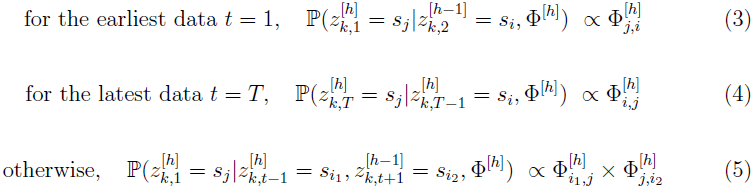

We performed the whole procedure *R* = 100 times. We ignored the first *B* = 100 “burn-in” iterations, leaving *R* × (*H − B*) = 4000 transition matrices for each ecoregion. The standard errors of the mean of the transition probabilities were consistently small, so we were finally able to derive the matrices of Figure 7 in main text (as well as Figures 2 and 3 in this appendix) as the mean values.

### 2 Shade tolerance classification across the US

**Table 1:** Shade tolerance indicators summarized for each US province.

| Province | Plots | Resampled plots | Distribution width | Spectral gap |
| --- | --- | --- | --- | --- |
| 232 | 91856 | 60751 | 0.5 | 0.069 |
| 212 | 82401 | 36975 | **0.9** | **0.023** |
| 231 | 54649 | 35165 | 0.6 | 0.081 |
| M221 | 23137 | 13696 | 0.7 | 0.060 |
| 223 | 24766 | 10578 | **0.8** | **0.035** |
| 221 | 20646 | 9491 | **0.8** | **0.035** |
| 222 | 23594 | 9462 | **0.9** | **0.033** |
| 211 | 14060 | 6003 | **0.8** | **0.034** |
| M211 | 10345 | 4712 | 0.6 | **0.047** |
| 251 | 9855 | 3478 | **0.8** | **0.040** |
| 234 | 2605 | 1062 | **0.8** | 0.064 |
| M231 | 2040 | 980 | 0.7 | 0.059 |
| M223 | 1268 | 677 | 0.7 | **0.047** |
| M331 | 5470 | 616 | 0.7 | **0.032** |
| 255 | 2846 | 542 | 0.7 | 0.085 |
| 332 | 1565 | 479 | 0.7 | **0.052** |
| 313 | 3499 | 452 | 0.4 | 0.058 |
| M261 | 4755 | 374 | 0.7 | **0.049** |
| 341 | 2338 | 353 | 0.4 | **0.046** |
| 411 | 585 | 328 | 0.7 | 0.113 |
| M341 | 2472 | 313 | 0.7 | 0.054 |
| 331 | 1819 | 295 | 0.7 | **0.043** |
| M334 | 634 | 295 | 0.5 | 0.073 |
| M242 | 5148 | 224 | **0.9** | 0.094 |
| M313 | 2130 | 174 | 0.5 | **0.052** |
| M332 | 4416 | 146 | 0.7 | NA |
| 321 | 1509 | 91 | 0.4 | NA |
| 322 | 470 | 65 | 0.4 | NA |
| 342 | 927 | 42 | 0.6 | NA |
| 263 | 418 | 34 | 0.6 | NA |
| M333 | 3105 | 33 | **0.9** | NA |
| M262 | 250 | 13 | 0.7 | NA |
| 242 | 544 | 13 | **0.8** | NA |
| 261 | 144 | 7 | **0.8** | NA |
| 262 | 5 | 0 | 0.5 | NA |
| 315 | 3597 | 0 | **0.8** | NA |

### 3 Dynamics of relative shade tolerance index

To investigate the dynamics of shade tolerance index in the early years of each area delimited in Section 3.2, we first computed for each ecoregion the value of shade relatively to their first value (that is, the average value of shade tolerance index for stands that are one year old), and pooled together ecoregions belonging to the same area (with either “wide” or “narrow” distribution, and either “fast” or “slow” convergence). We further modeled the dynamics with a piecewise linear model with two segments, by fitting them to the formula:

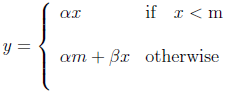

As we aligned the shade tolerance index to be 0 for the earliest stand age, this formula constraints the starting value *y* = 0. We performed the optimization using the Gauss-Newton algorithm, initialized with reasonable starting guesses (Bates and Watts, 1988). After convergence, all parameters *α*, m and *β* were found to be significantly different from 0 (one-sample two-sided t-tests, *n* between 7753 and 181718, *p <* 0.001). Due to the spread of the shade tolerance index, the fits were overall poor, with residual standard errors between and 0.135 and 0.260: while the established piecewise linear models are useful to highlight trends in the dataset, they should be used with caution when deriving quantitative predictions of shade tolerance index from the stand age.

The conceptual model predicts a decrease of the first stages followed by an increase (Figure 2 in main text). Specifically, ecoregions with a wide distribution of shade tolerance index exhibit the expected pattern regardless of their convergence speeds (bottom panels of Figure 1). Their computed inflection point, around 16-18 years, is further consistent with the study of the White Pine-Eastern Hemlock forests in Section 2.4. Ecoregions characterized as having a narrow distribution and a slow convergence exhibits an early decrease of shade tolerance, however there is barely any following increase (top-right panel of Figure 1, the *β* slope is one order of magnitude lower than the other parameters). Finally, the area characterized by a narrow distribution and a fast convergence exhibit a pattern exactly opposite to what we expect in the shade tolerance driven succession, with an early increase followed by a decrease.

**Figure 1:**
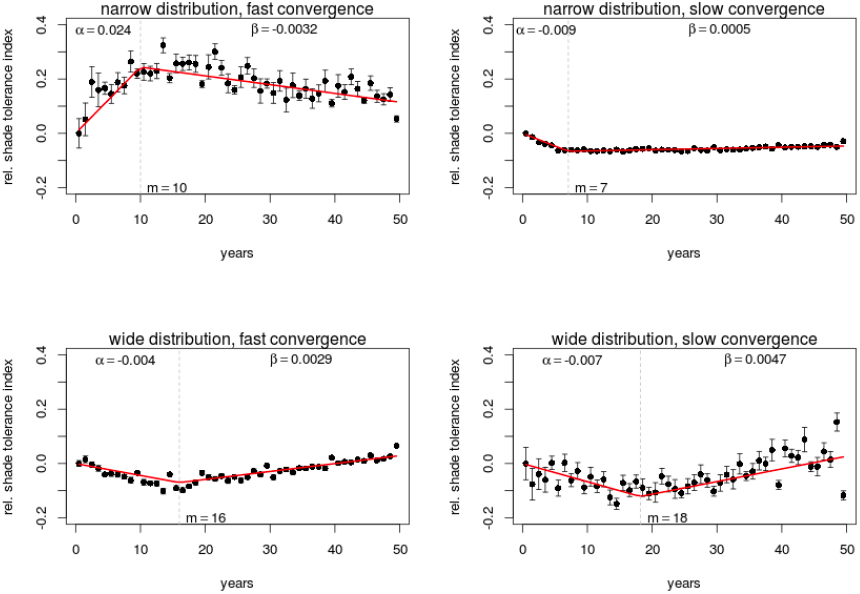
Relative shade tolerance index per area (in black, the error bars are standard errors of the mean) and piecewise linear regression (in red) with α being the slope of the first segment, β the slope of the second segment and m the age of the inflection point.

### 4 Transitions matrices computed on data subsets

**Figure 2:**
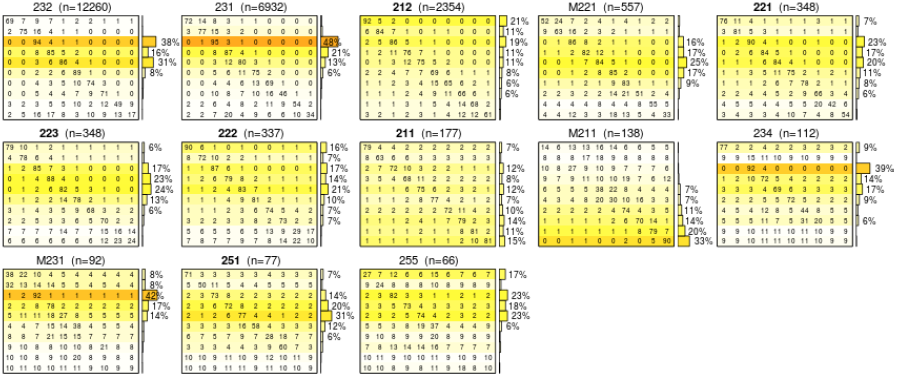
3-year transition matrices computed on the subset of plots with a stand age less than 20 years. The transitions are obviously much more noisy than the ones obtained on the full dataset (Figure 7 in main text), which is expected due to the very limited number of re-sampled young plots. Except for ecoregion “M211”, the early decrease of shade tolerance index predicted in the conceptual model is apparent in the matrices, with probabilities of decrease (transitions under the diagonal) superior to the probability of increase (transitions above the diagonal). The notations are similar to the ones in Figure 7 in main text.

**Figure 3:**
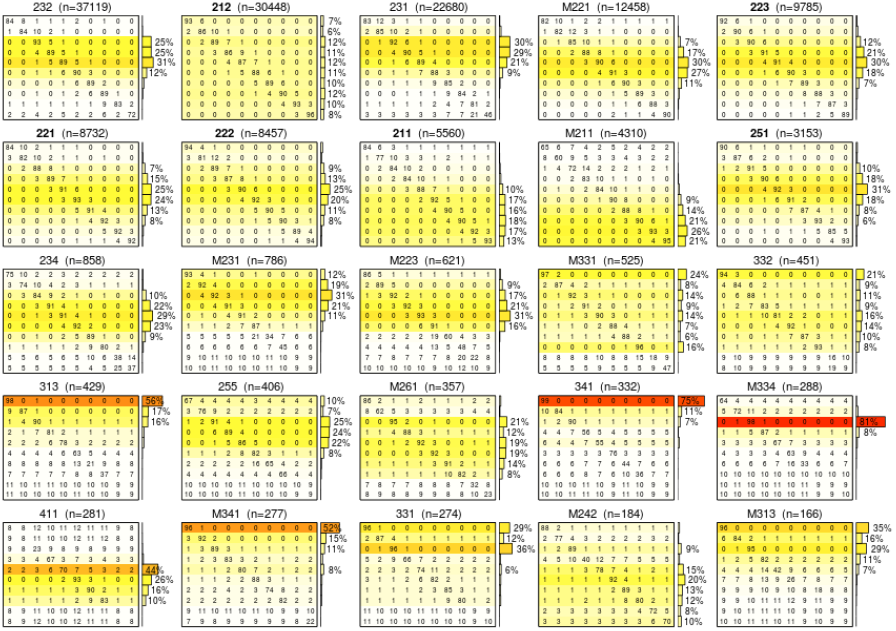
3-year transition matrices computed on the subset of plots with a stand age more or equal to 20 years. The transitions mostly match the ones obtained on the full dataset (Figure 7 in main text), confirming that transitions in the early stages (Figure 2 in this appendix) are not overall substantially contributing to the transition matrices. The notations are similar to the ones in Figure 7 in main text.

